# Bioelectrochemical engineering analysis of formate-mediated microbial electrosynthesis

**DOI:** 10.1101/2020.08.31.275537

**Authors:** Anthony J. Abel, Douglas S. Clark

**Author notes:** Correspondence should be addressed to D.S.C.

## Abstract

Mediated microbial electrosynthesis (MES) represents a promising strategy for the capture and conversion of CO_2_ into carbon-based products. We describe the development and application of a comprehensive multiphysics model to analyze a formate-mediated MES reactor. The model shows that this system can achieve a biomass productivity of ∼1.7 g L^-1^ hr^-1^ but is limited by a competitive trade-off between O_2_ gas/liquid mass transfer and CO_2_ transport to the cathode. Synthetic metabolic strategies are evaluated for formatotrophic growth, which can enable an energy efficiency of ∼21%, a 30% improvement over the Calvin cycle. However, carbon utilization efficiency is only ∼10% in the best cases due to a futile CO_2_ cycle, so gas recycle will be necessary for greater efficiency. Finally, separating electrochemical and microbial processes into separate reactors enables a higher biomass productivity of ∼2.4 g L^-1^ hr^-1^. The mediated MES model and analysis presented here can guide process design for conversion of CO_2_ into renewable chemical feedstocks.

## 1. Introduction

The capture and conversion of CO_2_ is a promising strategy for the production of carbon-based chemicals and could help to close the anthropogenic carbon cycle.^[1]^ Among the many strategies to fix CO_2_ using renewable energy, so-called “mediated” or “coupled” microbial electrosynthesis (MES) has received significant attention.^[2–5]^ In this scheme, electrons (ideally from a renewable source) are used to electrochemically reduce a mediator molecule that is then oxidized by planktonic microbes as a growth substrate.

Several groups have developed prototypical systems for mediated MES, relying on various redox mediators including H_2_,^[6–9]^ inorganic ions (*e*.*g*. ferrous ions or ammonia),^[10,11]^ simple organic molecules (*e*.*g*. carbon monoxide, formate, and methanol),^[12–15]^ and complex organic molecules such as the dye neutral red.^[16,17]^ Although these prototype systems have been able to achieve high efficiencies (∼10% in the case of Wang *et al*.^[8]^), the scalability and potential productivity of these systems remain unclear. The choice of redox mediator requires careful consideration: the ideal one should be abundantly available or easily produced electrochemically (eliminating complex organic molecules and inorganic ions), electropositive enough to directly reduce NAD(P)H for efficient energy transfer to cellular metabolism, and highly soluble in liquid water (eliminating H_2_).^[3,5]^ Formate/ic acid stands out as an especially promising redox mediator because it is readily and specifically produced from CO_2_^[18–20]^ and multiple natural and engineered formatotrophic growth mechanisms exist in workhorse bacteria.^[21–25]^

Initial scale-up,^[26]^ component integration,^[27]^ and media optimization^[28]^ studies have been performed for MES systems, demonstrating the need for careful attention to process parameters including the gas/liquid mass transfer coefficient (k_L_a). However, progress towards scaled, optimized systems has been limited, in part due to the complex nature of coupled bio-electrochemical systems. Because many physical processes occur simultaneously (diffusion and migration of species in fluid boundary layers, electrochemical and acid/base reactions, microbial growth and consumption and production of species, gas/liquid mass transfer, *etc*.), understanding the impact or potential of a given process or engineering strategy is difficult without a detailed, comprehensive model that accounts for all of the relevant physics. Moreover, such a model is necessary to quantify design and operation strategies that optimize efficiency and to identify process parameters that limit productivity.

To that end, several models of MES or related systems have been developed.^[29–34]^ Picioreanu *et al*. developed models for microbial fuel cells with planktonic microbes and biofilms using a generic redox mediator to shuttle electrons between the anode and microbes, both of which were able to accurately fit data from fed-batch experimental systems.^[32,33]^ Kazemi *et al*. developed one of the first MES models based on a conductive biofilm performing direct electron transfer, and using the model were able to relate applied potential to acetate production for the acetogenic bacterium *Sporomusa ovata* in a batch system.^[31]^ Recently, Gadkari *et al*. modeled microbial communities driving electrode reactions at both the anode and the cathode using a generic redox mediator, and predicted limiting production rates as a function of initial substrate concentration in a fed-batch system.^[29]^ Despite these successes, modeling studies have so far not focused on mediated MES systems or continuous operation schemes that will be necessary for processing at industrial scales. Moreover, previous models have focused on batch or fed-batch systems without considering physical phenomena such as gas/liquid mass transfer that are critical to scaled-up operation. Additionally, formate/ic acid requires explicit attention as a redox mediator because of its potential toxicity and participation in acid/base reactions that must be considered in reactor design and operation.

In this paper, we present a comprehensive multiphysics modeling framework that describes mass transport; electrochemical, acid/base, and microbial reaction kinetics; temperature effects, and gas/liquid mass transfer for an MES system generating formate and H_2_ (as a secondary product) at an abiotic cathode for consumption by planktonic cells (Fig. 1). The model is used to evaluate the effects of reactor design and operating parameters on critical performance metrics including biomass productivity, cell density, and carbon utilization and energy efficiency; and to compare the performance of microbes using different formatotrophic growth strategies under different optimal growth conditions. A key finding is that for integrated systems the tradeoff between O_2_ availability for microbial respiration and CO_2_ transport to the cathode surface for electrochemical reduction limits formate-mediated MES productivity and that separating electrochemical and microbial processes into two reactors avoids this fundamental limitation. The presented model, methodology, and analysis provide a complete framework for analyzing mediated MES reactor systems and identify promising research strategies for scale-up and process optimization that can advance MES systems from basic science to technological practice.

**Figure 1.**
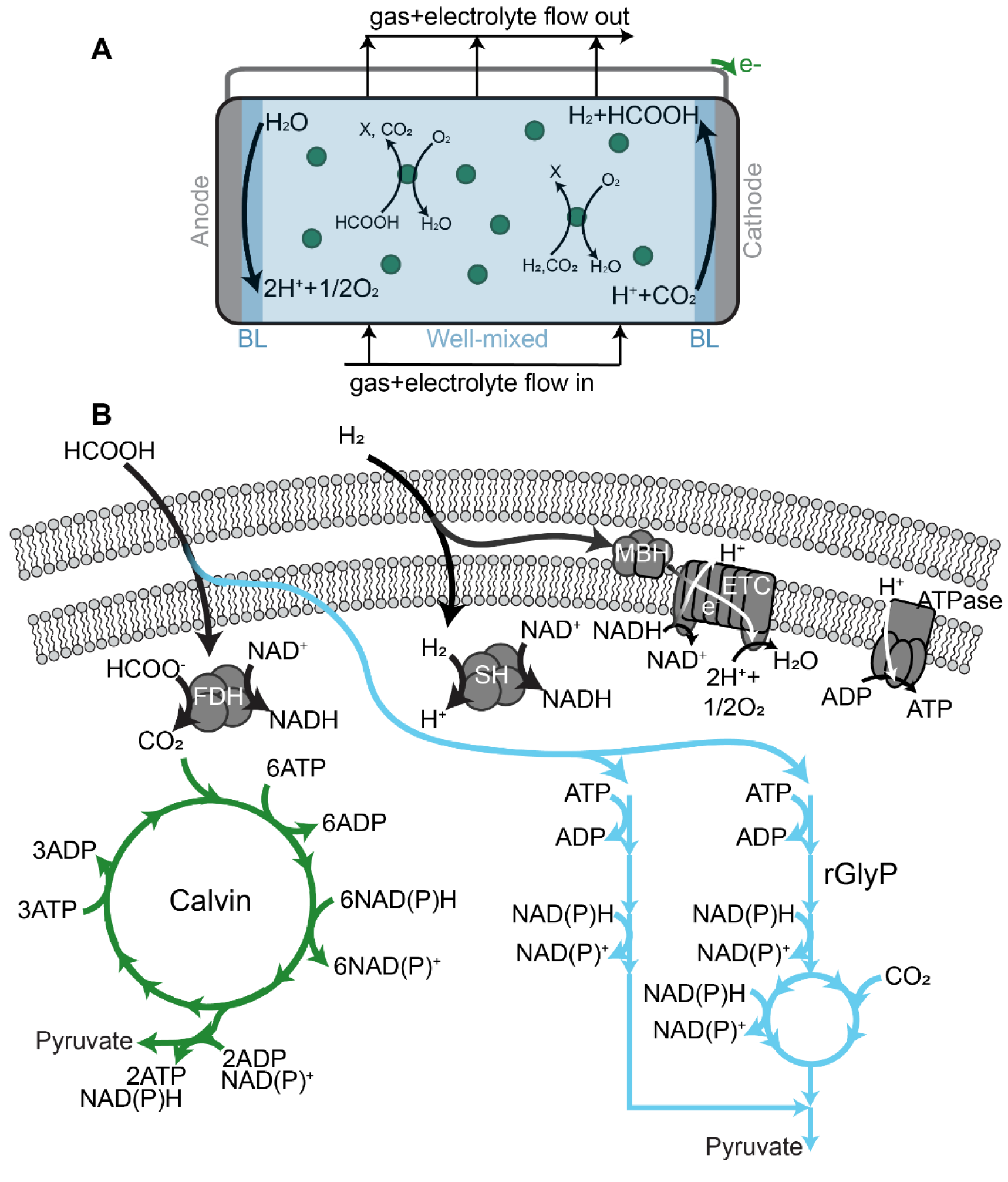
Reactor overview and formatotrophic growth strategies. (**A**) Reactor scheme. Carbon dioxide (CO_2_) and oxygen (O_2_) gas and electrolyte media are fed into a well-mixed bioelectrochemical reactor with hydrodynamic boundary layers separating the well-mixed phase from the electrode surfaces. (**B**) Energy metabolism (black, gray) and carbon fixation pathways (Calvin cycle: green, reductive glycine pathway (rGlyP): blue). X: biomass; FDH: formate dehydrogenase; SH: soluble hydrogenase; MBH: membrane-bound hydrogenase; ETC: electron transport chain.

## 2. Computational Methods

### 2.1 System overview and governing equations

The model considers a one-dimensional bio-electrochemical reactor for conversion of CO_2_ into biomass via formate (Fig. 1A). The reactor has a well-mixed region that is exchanged at a fixed dilution rate and to which a CO_2_/O_2_ gas mix is constantly supplied at a fixed pressure. Fluid boundary layers (BLs) separate the well-mixed liquid phase from the anode and cathode surfaces, where electrochemical reactions are driven by an applied voltage to oxidize water (at the anode surface) and reduce CO_2_ to HCOO^-^ or reduce protons to H_2_ (both at the cathode surface). Microbes at an initial concentration of *C*_*X*,0_ grow in the well-mixed phase by consuming HCOO^-^, H^+^, H_2_, CO_2_, and O_2_. The chemical species we consider in the reactor system are dissolved CO_2_, dissolved O_2_, dissolved H_2_, bicarbonate anions (HCO_3_^-^), carbonate anions (CO_3_^2-^), formic acid (HCOOH), formate anions (HCOO^-^), protons (H^+^), hydroxide anions (OH^-^), sodium cations (Na^+^), nitrate anions (NO_3_^-^), and microbes (X). NO_3_^-^ was selected as a representative anion for sodium salt to avoid the use of chloride ions (Cl^-^), which are known to produce deleterious and toxic side reactions at the cathode surface in MES systems.^[28]^ By neglecting ammonium/a species, we have assumed that they are fed in excess to the system as NH_3_. We consider the growth of two different model organisms: *Cupriavidus necator*, a well-studied chemolithoautotrophic organism in MES systems that is capable of growth on both formate and H_2_ as energy sources, and *Escherichia coli*, the biotechnology workhorse bacterium that has been recently engineered to support formatotrophic growth.^[22]^

#### 2.1.1 Well-mixed phase balance equations

The well-mixed electrolyte regions are assumed to have sufficient convective mixing such that no concentration gradients are formed. Such an open, well-mixed system must satisfy mass conservation, given generally for our reactor model by

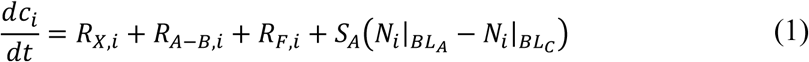

where *C*_*i*_ is the concentration, *R*_*i*_ is the net volumetric rate of formation and consumption due to microbial growth (X), acid/base reactions (A–B), and feed terms (F), and *N*_*i*_ is the flux of species *i*. The electrode surface area-to-volume ratio is given by *S*_*A*_. By convention, the positive x-direction is defined to the right of the page such that species flux from the cathode boundary layer phase (*BL*_*C*_) to the well-mixed phase will have a negative value.

#### 2.1.2 Species transport in the electrolyte boundary layers

The molar flux of species (assuming no net fluid velocity) in dilute electrolyte solutions is written as the sum of diffusive and migrative fluxes:

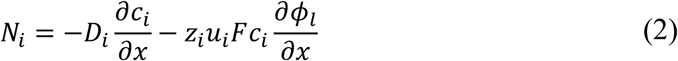

where *D*_*i*_ and *u*_*i*_ are the diffusivity and mobility (related by the Nernst-Einstein relationship, *u*_*i*_ = *D*_*i*_/*RT* for dilute solutions) and *z*_*i*_ is the charge number for species *i, F* is Faraday’s constant, and *ϕ*_*l*_ is the local electrolyte potential. Flux boundary conditions at the anode and cathode surfaces are described in section 2.4.

#### 2.1.3 Charge conservation and electroneutrality

Charge conservation in the system requires that

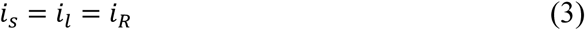

where *i*_*n*_ is the current density in the solid electrode (*s*), electrolyte media (*l*), and at the electrode/electrolyte interface (*R, i*.*e*. the electrochemical reaction current density). The net ionic current density (*i*_*l*_) can be calculated from the total ionic flux,

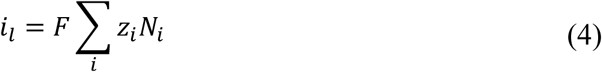

following electroneutrality, given by

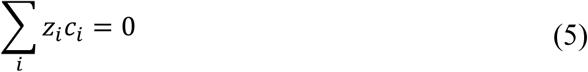

We note that the equilibrium distribution of species in gas-saturated electrolyte media is constrained by the electroneutrality requirement, the four independent acid/base reactions given in section 2.3, the three gas solubility relationships given in section 2.5, and the requirement that mole fractions sum to unity. Therefore, this system has two degrees of freedom (excluding microbes), and is fully constrained by setting the concentrations of, for example, CO_2_ and H^+^ (or the pH). To better replicate experimental procedures, we fix the pH and the NaNO_3_ concentration and use the equilibrium, solubility, and electroneutrality relationships to determine all other values.

### 2.2 Microbial growth

Microbial growth occurs in the well-mixed phase and is responsible for the production of more cells and the consumption or production of several chemical species. These reactions are compiled in *R*_*X,i*_, which is written as

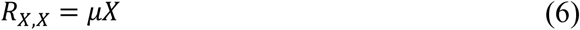

for microbes, and as:

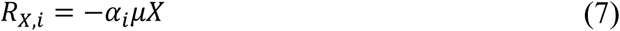

for all other species. Here, *μ* is the specific growth rate and *α*_*i*_ is a stoichiometric coefficient, which will be defined for different species as *α, β, γ, κ, ϵ*, and *ζ*, following convention from Blanch and Clark,^[35]^ in the following sections.

We define microbial growth kinetics using the Monod model. For formatotrophic growth, we consider the growth rate dependence on both formate and oxygen:

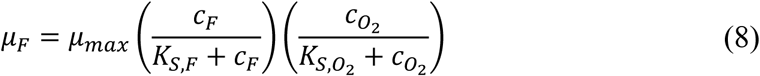

where *μ*_*F*_ refers to growth on formate, *μ*_*max*_ is the maximum specific growth rate, and *K*_*S,i*_ is the Monod constant or half-saturation constant for substrate *i*. For hydrogenotrophic growth, we consider the concentrations of H_2_, O_2_, CO_2_, and HCOO^-^ (only for growth using the reductive Glycine pathway) following the same method:

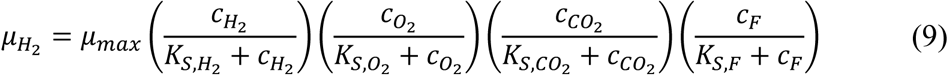

In the case of *C. necator* growth, we simply sum the growth on formate and H_2_:

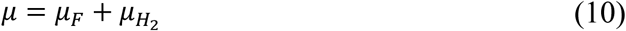

#### 2.2.1 Formatotrophic growth yield

We use a simple, generic equation to describe both complete formate oxidation coupled with CO_2_ fixation and partial formate oxidation coupled to partial formate assimilation (Fig. 1B):

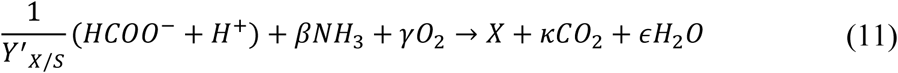

The molar cell yield, *Y*^′^_*X*/*S*_, is influenced by formate concentration due to a range of toxicity effects in *C. necator*, and is given by an empirical equation^[23]^:

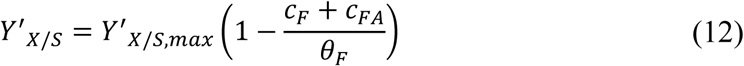

where *θ*_*F*_ is a fitting parameter that represents the maximum formate/ic acid concentration at which cells can grow. By mole balance, the stoichiometric coefficients are

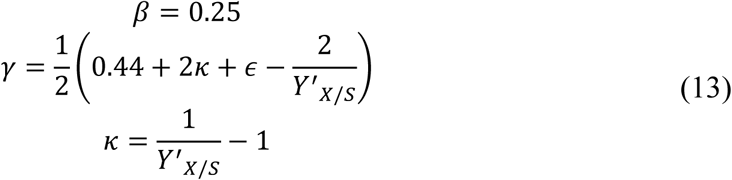

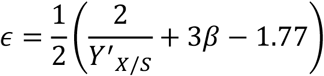

for biomass with the composition *C*_1_*H*_1.77_*O*_0.44_*N*_0.25_.^[23]^

The reductive glycine pathway (rGlyP), recently engineered in both *C. necator*^[21]^ and *E. coli*^[22]^ and discovered in wild-type phosphite-oxidizing organisms,^[36]^ is predicted to enable higher biomass yield using formate as a growth substrate than the Calvin cycle (Fig. 1B).^[24]^ To evaluate the promise of this alternate formatotrophic growth strategy, we modeled the improved growth by increasing *Y*^′^_*X*/*S,max*_ 27% relative to its value for the Calvin cycle based on the predicted theoretical improvement (see supplementary note A for additional details).

We used the same equations (11 – 13) to describe formatotrophic growth of *E. coli* using the rGlyP, and adjusted *θ*_*F*_ to reflect the maximum formate concentration that enabled growth reported by Kim *et al*.^[22]^ We used *Y*^′^_*X*/*S,max*_ and *μ*_*max*_ values equivalent to the theoretical values for *C. necator* using the rGlyP to reflect an optimistic outlook on the promise of further engineering to improve formatotrophic *E. coli* growth (the strain reported by Kim *et al*. achieves ∼42% of this yield value and ∼50% of the maximum growth rate).

#### 2.2.2 Hydrogenotrophic growth yield

A simple equation for hydrogenotrophic growth of *C. necator* using the Calvin cycle is given by Ishizaki *et al*.^[37]^:

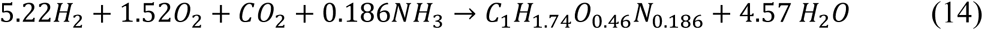

We note that the cell stoichiometry measured by Ishizaki *et al*. is slightly different from that used by Grunwald *et al*.^[23]^ In our model, cell mass is a single species that is not broken into its constitutive elements, so we don’t account for this difference in our system.

To compare hydrogenotrophic growth using the Calvin cycle and the rGlyP, we generalize eq. (14) according to

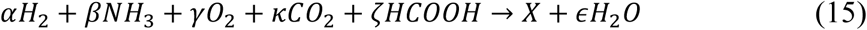

where *ζ* = 0 for growth with the Calvin cycle and *ζ* = 2*κ* for growth on the rGlyP (see supplementary note A for additional details). The stoichiometric coefficients are determined by a mole balance using the cell stoichiometry given by Ishizaki *et al*.^[37]^:

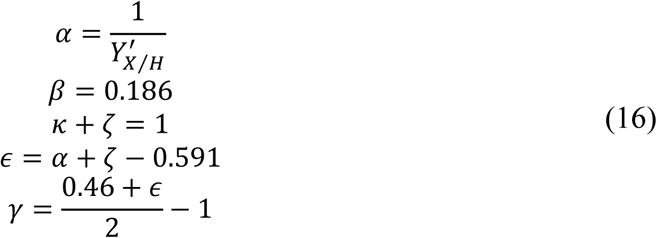

Because the rGlyP makes more efficient use of reducing equivalents, we increase the biomass yield 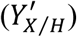 by ∼82% as predicted by the theoretical calculations (Supplementary note A).

#### 2.2.3 Growth rate dependence on temperature and pH

We use a simple model to describe the effects of temperature and pH on microbial growth following Rosso *et al*.^[38]^:

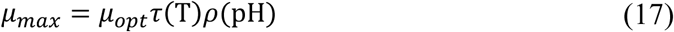

where *μ*_*opt*_ is the growth rate at optimal conditions and *τ*(T) and *ρ*(pH) are written as

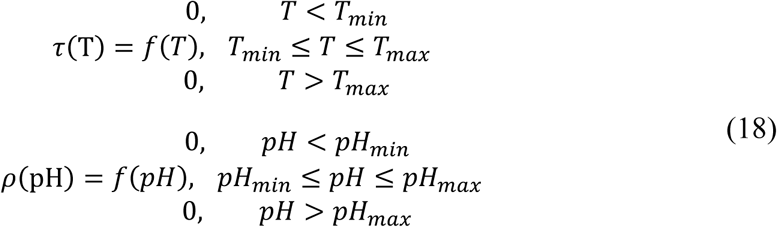

Here, *T*_*min*/*max*_ and *pH*_*min*/*max*_ are the ranges of temperature and pH over which microbial growth is observed, and the functions *f*(*T*) and *f*(*pH*) are

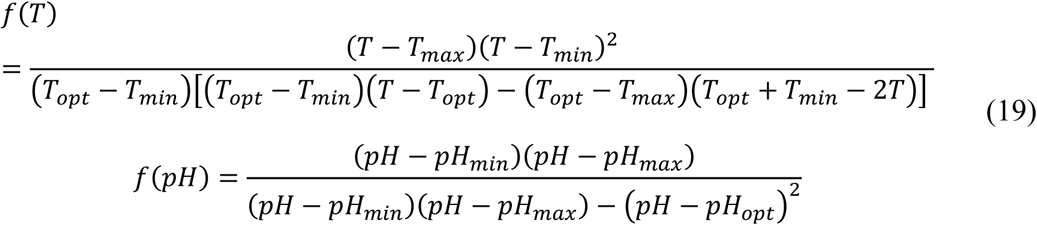

where *T*_*opt*_ and *pH*_*opt*_ are the optimal temperature and pH for growth, respectively.

### 2.3 Acid/base reactions

The acid/base bicarbonate/carbonate, formic acid/formate, and water dissociation reactions shown below occur in all phases and are treated as kinetic expressions without assuming equilibrium (eq. (26)):

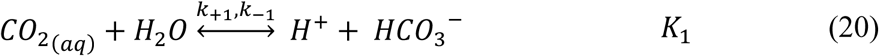

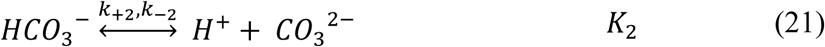

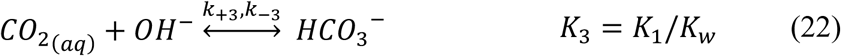

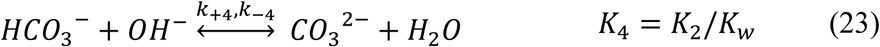

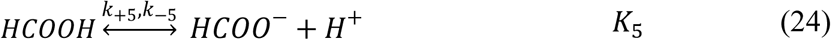

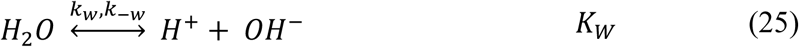

where *k*_+*n*_ and *k*_−*n*_ are the forward and reverse rate constants, respectively, and *K*_*n*_ is the equilibrium constant for the *n*th reaction, given by

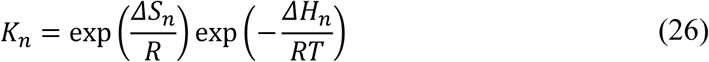

Source and sink terms resulting from these reactions are compiled in *R*_*A*−*B,i*_, written as

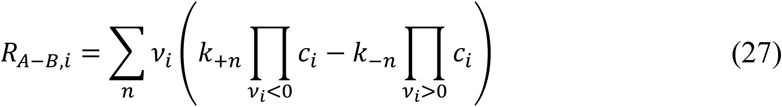

where *v*_*i*_ is the stoichiometric coefficient of species *i* for the *n*th reaction and reverse rate constants (*k*_−*n*_) are calculated from:

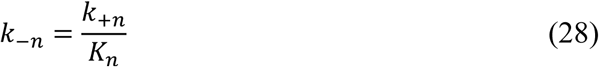

### 2.4 Electrochemical reactions and electron transport

The surface reaction at the anode is the oxidation of water:

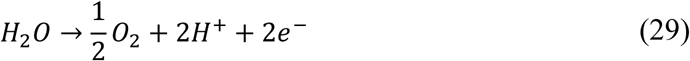

which we write in its acidic form to reflect the fact that acidic conditions are observed at the anode surface. At the cathode, we consider two reduction reactions:

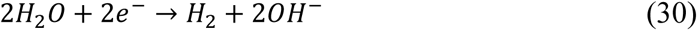

and

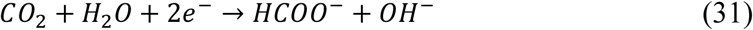

which are both described under the basic conditions observed at the cathode. The surface reactions relate current density to species generation, consumption, and transport by flux boundary conditions given by

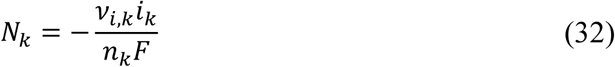

where *i*_*k*_ is the current density, *v*_*i,k*_ is the stoichiometric coefficient of species *i*, and *n*_*k*_ is the number of participating electrons for electrochemical reaction *k*.

#### 2.4.1 Electrochemical kinetics

Charge transfer reactions occur at the electrode/electrolyte interface and can be described by the Butler-Volmer equation:

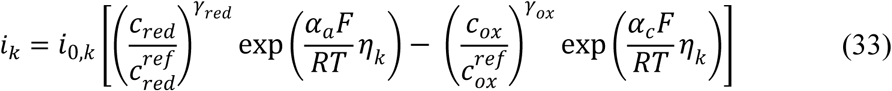

where *i*_0,*k*_ is the exchange current density for reaction *k, γ*_*red*/*ox*_ is the reaction order with respect to some reactant *C*_*red*/*ox*_, *α*_*a*/*C*_ is the anodic/cathodic transfer coefficient, and *η*_*k*_ is the overpotential for reaction *k*. The exchange current density, *i*_0,*k*_ depends on a pre-exponential factor (*A*_*k*_) and an apparent activation energy (*E*_*a,k*_) that can be pH-dependent according to the Arrhenius equation,

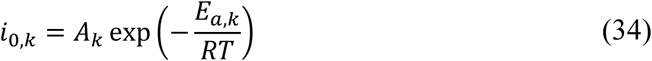

The overpotential, *η*_*k*_, is defined according to

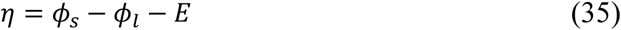

where *ϕ*_*s*_ is the electrode potential, *ϕ*_*l*_ is the electrolyte potential, and *E* is the half-cell equilibrium potential.

#### 2.4.2 Electron transport in the solid electrodes

Electron transport in the solid electrode regions is governed by charge conservation (eq. (3)) and Ohm’s law:

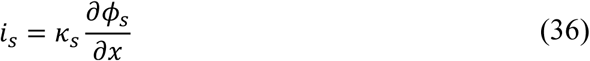

where *κ*_*s*_ is the anode or cathode conductivity.

### 2.5 Gas feed and electrolyte flow

#### 2.5.1 Gas feed

A CO_2_/O_2_ gas mixture at a pressure *P* is fed to the reactor, resulting in mass transfer into the liquid phase according to

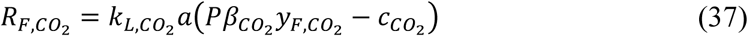

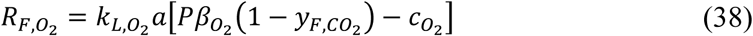

where *k*_*L,i*_*a* is the volumetric mass-transfer coefficient on the liquid side of the gas/liquid interface, *β*_*i*_ is the Bunsen solubility coefficient, and *y*_*F,i*_ is the mole fraction of species *i* in the gas phase.

#### 2.5.2 Equilibrium solubility of CO_2_, O_2_, and H_2_ in electrolyte

We calculate the equilibrium solubility of CO_2_, O_2_, and H_2_ according to the empirical relationship for the Bunsen solubility coefficient (*β*):

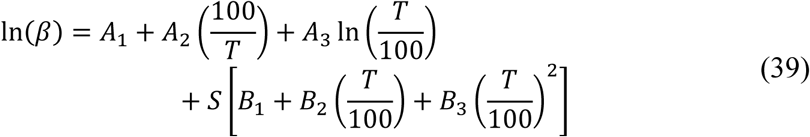

where *A*_*n*_ and *B*_*n*_ are fitting parameters and *S* is the electrolyte salinity in g/kg water. The equilibrium concentration of gaseous species in the liquid phase is then given simply by

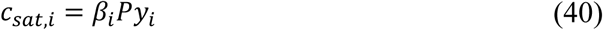

#### 2.5.3 Gas/liquid mass transfer coefficients

Gas/liquid mass transfer coefficients can be calculated either from first principles^[35]^ or by any of several correlations that are dependent on the system geometry. Here, we use the correlation first developed by Vasconcelos *et al*. for stirred tank reactors with a height that is twice the diameter^[39]^:

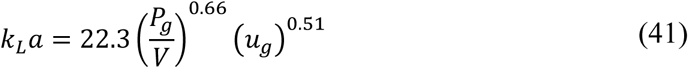

where *P*_*g*_/*V* is the specific power input (in units W m^-3^) and *u*_*g*_ is the superficial gas velocity (in units m s^-1^). In our model, we assume a *k*_*L*_*a* value of 300 hr^-1^, which corresponds to a power demand of 4000 W m^-3^ with a superficial gas velocity of ∼0.11 m s^-1^. We note that the actual *k*_*L*_*a* value in a given reactor is highly dependent on the gas feeding mechanism,^[40]^ the reactor geometry and gas contacting strategies,^[41]^ and components integrated into bioreactors,^[27]^ so any model of experimental results must rely on carefully measured *k*_*L*_*a* values before valid comparisons can be made.

#### 2.5.4 Evolution of supersaturated gas at electrode surfaces

Water oxidation at the anode surface and hydrogen evolution at the cathode surface will generate O_2_ and H_2_ in excess of what the liquid phase can solubilize. Additionally, because water oxidation creates acidic conditions near the anode surface, bicarbonate and carbonate species will be converted to aqueous CO_2_ according to Le Chatelier’s principle. To avoid the unrealistic supersaturation of gases in the electrolyte media these mechanisms would cause, we describe evolution of gases as a flux boundary condition at the electrode surfaces as:

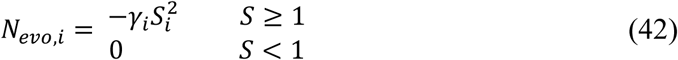

where *γ*_*i*_ is the releasing coefficient and *S*_*i*_ is the supersaturation coefficient for species *i*, defined as *C*_*i*_/*β*_*i*_*p*_*i*_, where *p*_*i*_ is the partial pressure. The releasing coefficient, *γ*_*i*_, is known to vary over at least seven orders of magnitude.^[42]^ Lin *et al*. studied the rate of CO_2_ evolution driven by electrolyte acidification in a CO_2_ electrolyzer system and showed that the value of the releasing coefficient does not impact the steady-state rate of gas evolution but does change the time to reach steady-state evolution conditions.^[43]^ In their system, gas evolution reached steady-state values in <30 min when the current density was ∼10 mA cm^-2^, so the actual value of *γ*_*i*_ is not expected to change the conclusions of our model.

#### 2.5.5 Gaseous species concentration exiting the reactor

Only a fraction of the CO_2_ (and O_2_) fed to the reactor is transferred to the liquid phase; defining this fraction as 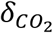 allows us to calculate the flow rate of fed gas to the reactor (*f*_*F*_), given by:

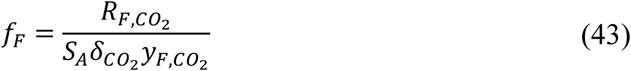

where *S*_*A*_ is the reactor electrode surface area to reactor volume ratio. Note that *f*_*F*_ is written in units of moles per area per time (mol l^-2^ t^-1^); for generality, we have normalized our feed rates to the electrode surface area to more straightforwardly connect this feed rate to the rates of gas evolution from the electrode surfaces. We can also define the fraction of fed O_2_ transferred to the liquid phase, 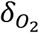, as

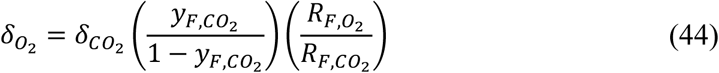

by performing a mass balance on the feed gas stream.

Gas exiting the reactor will include CO_2_, O_2_, and H_2_. To calculate their mole fractions, we perform a mole balance on the gas phase of the reactor and assume perfect mixing. The flow rate of gas exiting the reactor is given by:

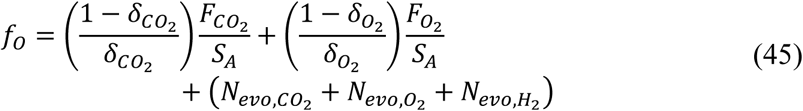

Here, the first two terms represent CO_2_ and O_2_ fed to the reactor that do not dissolve into the liquid phase, and the final term represents fluxes of gas species due to evolution at electrode surfaces, as described in eq. 42.

Gaseous species mole fractions of the exiting gas stream are then calculated by component mole balances:

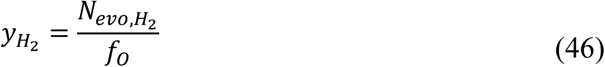

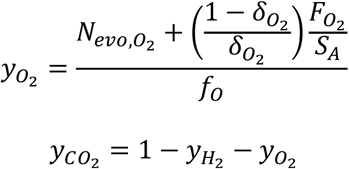

We use these equations to predict the saturation concentration of H_2_ in the liquid phase. They could also be used to identify the operational conditions that lead to flammable gas mixtures of H_2_ and O_2_ in the reactor headspace. In our model, 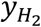 was ≤5% in all cases when using 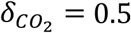, so flammable conditions are not expected to occur under typical operation.

#### 2.5.6 Electrolyte media flow

Electrolyte media is fed to and extracted from the well-mixed liquid phase at a constant dilution rate, resulting in a feed term written as

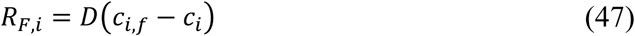

where *D* is the dilution rate (defined as the inverse space time, or volumetric flow rate divided by reactor volume^[35]^). We assume the feed stream is free of microbes but otherwise equivalent to the initial conditions (*i*.*e. C*_*i*≠*X,f*_ = *C*_*i*≠*X*,0_; *C*_*X,f*_ = 0) and that CO_2_ and O_2_ are supplied by the gas feed.

### 2.6 Model implementation

The governing equations are solved using the MUMPS general solver in COMSOL Multiphysics 5.4 with a nonlinear controller. The modeling domain has a maximum element size of 20 μm in the well-mixed regions and 2 μm in boundary layers to capture concentration gradients; the solution was independent of increasing mesh resolution. Model parameters are listed in Table 1. The potential in the reactor is calculated relative to zero potential at the cathode base and potential is applied as a boundary condition at the anode.

**Table 1.**
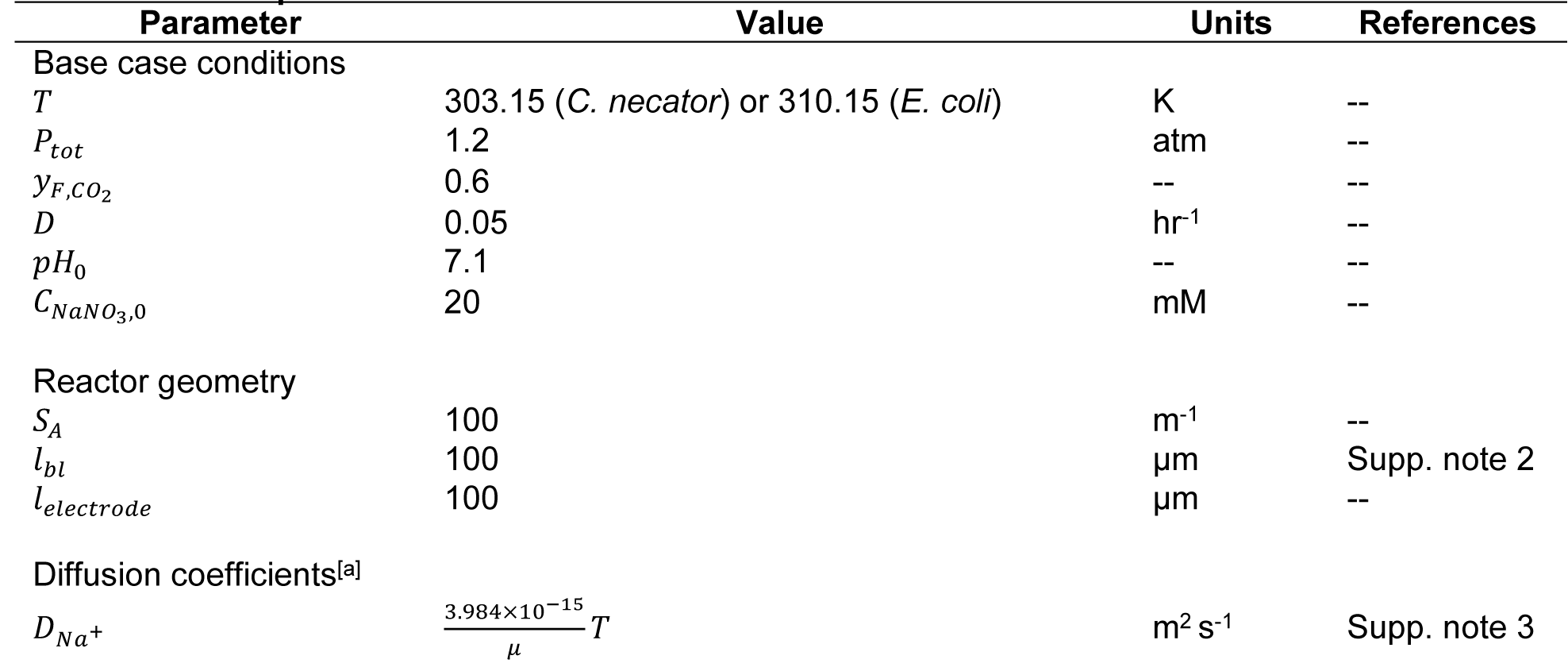
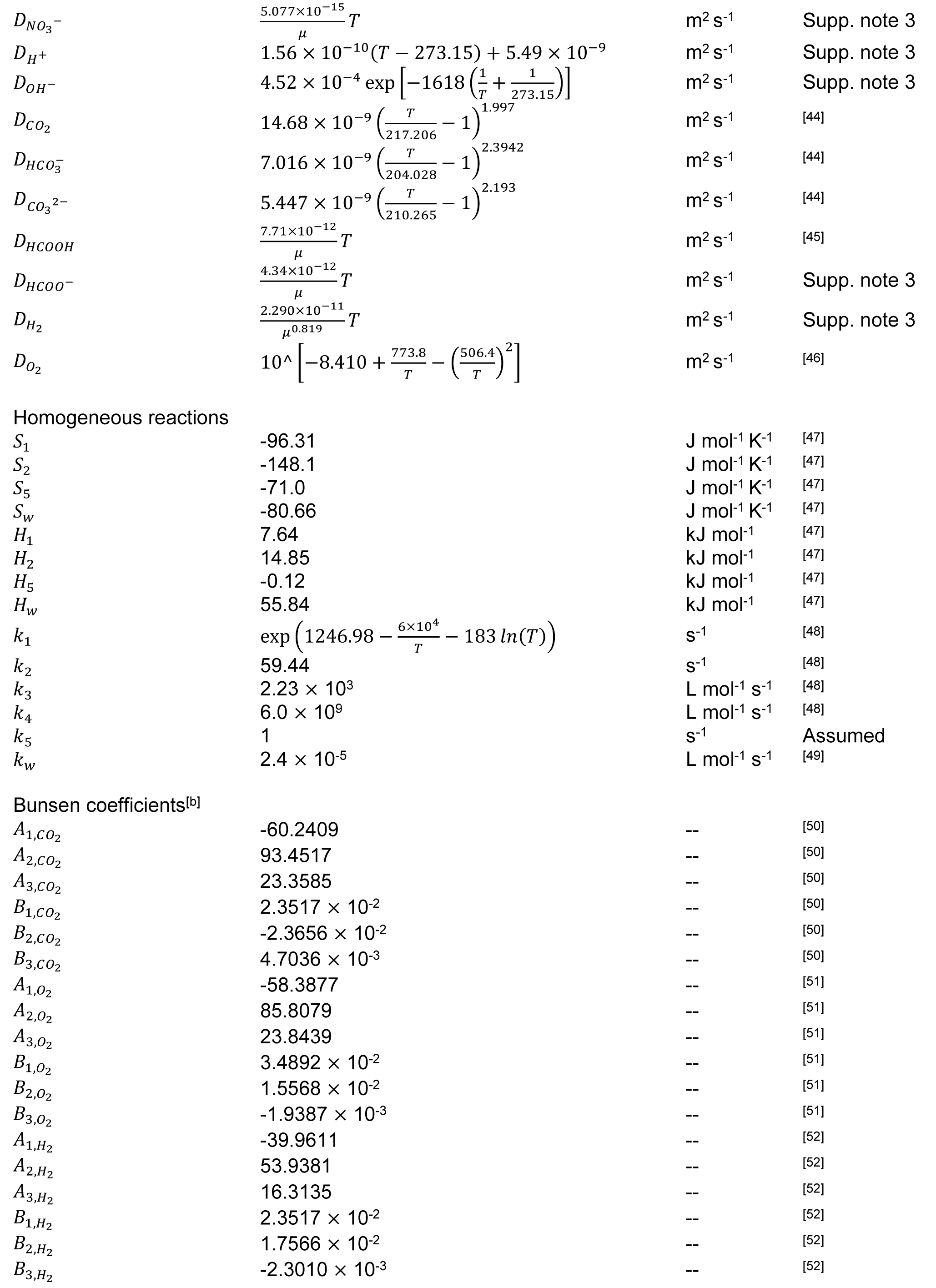
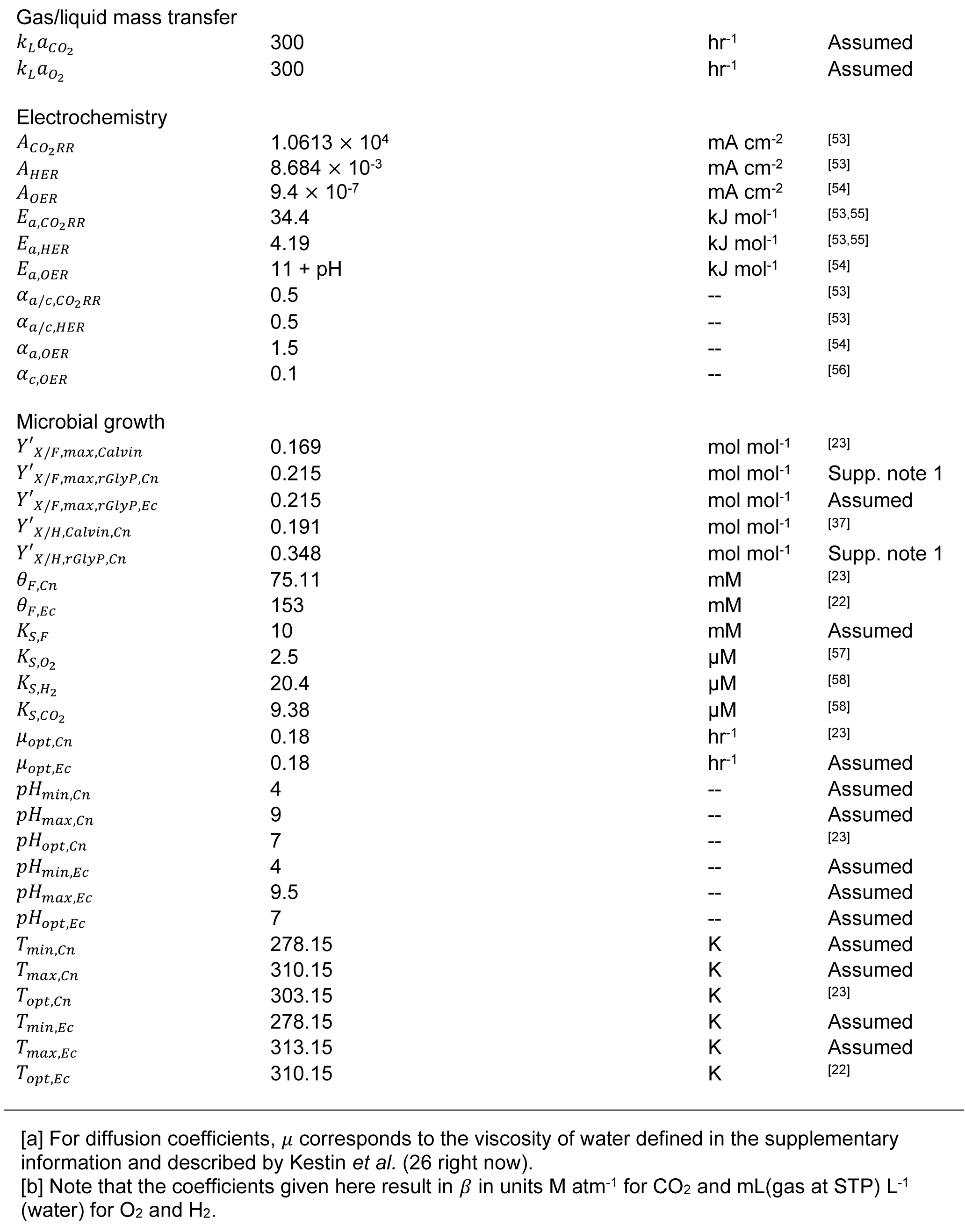
Model parameters.

| Parameter | Value | Units | References |
| --- | --- | --- | --- |
| Base case conditions |  |  |  |
| $T$ | 303.15 (*C. necator*) or 310.15 (*E. coli*) | K | -- |
| $P_{tot}$ | 1.2 | atm | -- |
| $y_{F,CO_2}$ | 0.6 | -- | -- |
| $D$ | 0.05 | hr<sup>-1</sup> | -- |
| $pH_0$ | 7.1 | -- | -- |
| $C_{NaNO_3,0}$ | 20 | mM | -- |
| Reactor geometry |  |  |  |
| $S_A$ | 100 | m<sup>-1</sup> | -- |
| $l_{bl}$ | 100 | $\mu m$ | Supp. note 2 |
| $l_{electrode}$ | 100 | $\mu m$ | -- |
| Diffusion coefficients<sup>[a]</sup> |  |  |  |
| $D_{Na^+}$ | $\frac{3.984 \times 10^{-15}}{\mu} T$ | m<sup>2</sup> s<sup>-1</sup> | Supp. note 3 |
| $D_{NO_3^-}$ | $\frac{5.077 \times 10^{-15}}{\mu} T$ | $m^2 s^{-1}$ | Supp. note 3 |
| $D_{H^+}$ | $1.56 \times 10^{-10} (T - 273.15) + 5.49 \times 10^{-9}$ | $m^2 s^{-1}$ | Supp. note 3 |
| $D_{OH^-}$ | $4.52 \times 10^{-4} \exp \left[ -1618 \left( \frac{1}{T} + \frac{1}{273.15} \right) \right]$ | $m^2 s^{-1}$ | Supp. note 3 |
| $D_{CO_2}$ | $14.68 \times 10^{-9} \left( \frac{T}{217.206} - 1 \right)^{1.997}$ | $m^2 s^{-1}$ | [44] |
| $D_{HCO_3^-}$ | $7.016 \times 10^{-9} \left( \frac{T}{204.028} - 1 \right)^{2.3942}$ | $m^2 s^{-1}$ | [44] |
| $D_{CO_3^{2-}}$ | $5.447 \times 10^{-9} \left( \frac{T}{210.265} - 1 \right)^{2.193}$ | $m^2 s^{-1}$ | [44] |
| $D_{HCOOH}$ | $\frac{7.71 \times 10^{-12}}{\mu} T$ | $m^2 s^{-1}$ | [45] |
| $D_{HCOO^-}$ | $\frac{4.34 \times 10^{-12}}{\mu} T$ | $m^2 s^{-1}$ | Supp. note 3 |
| $D_{H_2}$ | $\frac{2.290 \times 10^{-11}}{\mu^{0.819}} T$ | $m^2 s^{-1}$ | Supp. note 3 |
| $D_{O_2}$ | $10^{\left[ -8.410 + \frac{773.8}{T} - \left( \frac{506.4}{T} \right)^2 \right]}$ | $m^2 s^{-1}$ | [46] |
| Homogeneous reactions |  |  |  |
| $S_1$ | -96.31 | $J mol^{-1} K^{-1}$ | [47] |
| $S_2$ | -148.1 | $J mol^{-1} K^{-1}$ | [47] |
| $S_5$ | -71.0 | $J mol^{-1} K^{-1}$ | [47] |
| $S_w$ | -80.66 | $J mol^{-1} K^{-1}$ | [47] |
| $H_1$ | 7.64 | $kJ mol^{-1}$ | [47] |
| $H_2$ | 14.85 | $kJ mol^{-1}$ | [47] |
| $H_5$ | -0.12 | $kJ mol^{-1}$ | [47] |
| $H_w$ | 55.84 | $kJ mol^{-1}$ | [47] |
| $k_1$ | $\exp \left( 1246.98 - \frac{6 \times 10^4}{T} - 183 \ln(T) \right)$ | $s^{-1}$ | [48] |
| $k_2$ | 59.44 | $s^{-1}$ | [48] |
| $k_3$ | $2.23 \times 10^3$ | $L mol^{-1} s^{-1}$ | [48] |
| $k_4$ | $6.0 \times 10^9$ | $L mol^{-1} s^{-1}$ | [48] |
| $k_5$ | 1 | $s^{-1}$ | Assumed |
| $k_w$ | $2.4 \times 10^{-5}$ | $L mol^{-1} s^{-1}$ | [49] |
| Bunsen coefficients<sup>[b]</sup> |  |  |  |
| $A_{1,CO_2}$ | -60.2409 | -- | [50] |
| $A_{2,CO_2}$ | 93.4517 | -- | [50] |
| $A_{3,CO_2}$ | 23.3585 | -- | [50] |
| $B_{1,CO_2}$ | $2.3517 \times 10^{-2}$ | -- | [50] |
| $B_{2,CO_2}$ | $-2.3656 \times 10^{-2}$ | -- | [50] |
| $B_{3,CO_2}$ | $4.7036 \times 10^{-3}$ | -- | [50] |
| $A_{1,O_2}$ | -58.3877 | -- | [51] |
| $A_{2,O_2}$ | 85.8079 | -- | [51] |
| $A_{3,O_2}$ | 23.8439 | -- | [51] |
| $B_{1,O_2}$ | $3.4892 \times 10^{-2}$ | -- | [51] |
| $B_{2,O_2}$ | $1.5568 \times 10^{-2}$ | -- | [51] |
| $B_{3,O_2}$ | $-1.9387 \times 10^{-3}$ | -- | [51] |
| $A_{1,H_2}$ | -39.9611 | -- | [52] |
| $A_{2,H_2}$ | 53.9381 | -- | [52] |
| $A_{3,H_2}$ | 16.3135 | -- | [52] |
| $B_{1,H_2}$ | $2.3517 \times 10^{-2}$ | -- | [52] |
| $B_{2,H_2}$ | $1.7566 \times 10^{-2}$ | -- | [52] |
| $B_{3,H_2}$ | $-2.3010 \times 10^{-3}$ | -- | [52] |

|  |  |  |  |  |
| --- | --- | --- | --- | --- |
| Gas/liquid mass transfer |  |  |  |  |
| $k_L a_{CO_2}$ | 300 | hr<sup>-1</sup> | Assumed | |
| $k_L a_{O_2}$ | 300 | hr<sup>-1</sup> | Assumed | |
| Electrochemistry |  |  |  |  |
| $A_{CO_2RR}$ | $1.0613 \times 10^4$ | mA cm<sup>-2</sup> | [53] | |
| $A_{HER}$ | $8.684 \times 10^{-3}$ | mA cm<sup>-2</sup> | [53] | |
| $A_{OER}$ | $9.4 \times 10^{-7}$ | mA cm<sup>-2</sup> | [54] | |
| $E_{a,CO_2RR}$ | 34.4 | kJ mol<sup>-1</sup> | [53,55] | |
| $E_{a,HER}$ | 4.19 | kJ mol<sup>-1</sup> | [53,55] | |
| $E_{a,OER}$ | 11 + pH | kJ mol<sup>-1</sup> | [54] | |
| $\alpha_{a/c,CO_2RR}$ | 0.5 | -- | [53] | |
| $\alpha_{a/c,HER}$ | 0.5 | -- | [53] | |
| $\alpha_{a,OER}$ | 1.5 | -- | [54] | |
| $\alpha_{c,OER}$ | 0.1 | -- | [56] | |
| Microbial growth |  |  |  |  |
| $Y'_{X/F,max,Calvin}$ | 0.169 | mol mol<sup>-1</sup> | [23] | |
| $Y'_{X/F,max,rGlyP,Cn}$ | 0.215 | mol mol<sup>-1</sup> | Supp. note 1 | |
| $Y'_{X/F,max,rGlyP,Ec}$ | 0.215 | mol mol<sup>-1</sup> | Assumed | |
| $Y'_{X/H,Calvin,Cn}$ | 0.191 | mol mol<sup>-1</sup> | [37] | |
| $Y'_{X/H,rGlyP,Cn}$ | 0.348 | mol mol<sup>-1</sup> | Supp. note 1 | |
| $\theta_{F,Cn}$ | 75.11 | mM | [23] | |
| $\theta_{F,Ec}$ | 153 | mM | [22] | |
| $K_{S,F}$ | 10 | mM | Assumed | |
| $K_{S,O_2}$ | 2.5 | μM | [57] | |
| $K_{S,H_2}$ | 20.4 | μM | [58] | |
| $K_{S,CO_2}$ | 9.38 | μM | [58] | |
| $\mu_{opt,Cn}$ | 0.18 | hr<sup>-1</sup> | [23] | |
| $\mu_{opt,Ec}$ | 0.18 | hr<sup>-1</sup> | Assumed | |
| $pH_{min,Cn}$ | 4 | -- | Assumed | |
| $pH_{max,Cn}$ | 9 | -- | Assumed | |
| $pH_{opt,Cn}$ | 7 | -- | [23] | |
| $pH_{min,Ec}$ | 4 | -- | Assumed | |
| $pH_{max,Ec}$ | 9.5 | -- | Assumed | |
| $pH_{opt,Ec}$ | 7 | -- | Assumed | |
| $T_{min,Cn}$ | 278.15 | K | Assumed | |
| $T_{max,Cn}$ | 310.15 | K | Assumed | |
| $T_{opt,Cn}$ | 303.15 | K | [23] | |
| $T_{min,Ec}$ | 278.15 | K | Assumed | |
| $T_{max,Ec}$ | 313.15 | K | Assumed | |
| $T_{opt,Ec}$ | 310.15 | K | [22] | |

[a] For diffusion coefficients, $\mu$ corresponds to the viscosity of water defined in the supplementary information and described by Kestin *et al.* (26 right now).
[b] Note that the coefficients given here result in $\beta$ in units M atm<sup>-1</sup> for CO<sub>2</sub> and mL(gas at STP) L<sup>-1</sup> (water) for O<sub>2</sub> and H<sub>2</sub>.

## 3. Results and Discussion

### 3.1 CO_2_ diffusion and O_2_ interphase transfer determine upper bounds on productivity

We first evaluated the performance of a mediated MES system using *C. necator* as the microbial catalyst since a wealth of formatotrophic and hydrogenotrophic growth data for this organism exists in the literature (Fig. 2).^[21,23,37]^ We used experimental parameters describing electrochemical reduction towards HCOO^-^ and H_2_ on a Sn electrode because of its ability to selectively produce formate^[18,53]^ and we compared operation with two different gas feed compositions: 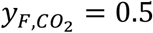, corresponding to an equal mixture of CO_2_ and O_2_ in the gas feed stream; and 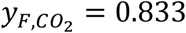, corresponding to a CO_2_ partial pressure of 1 atm for our model. Current density towards CO_2_ reduction or H_2_ production increases exponentially as a function of voltage following Butler-Volmer kinetics (Fig. 2A, eq. (33)). Because the cell density and biomass productivity depend linearly on the current density (see eq. S25, for example), these values also increase exponentially with the applied voltage (Fig. 2B, C). For the equimolar gas feed mixture (dashed curves in Fig. 2), the molar yield remains constant at just below its maximum value, indicating that microbes are consuming nearly all the formate produced by the electrochemical reaction (*i*.*e*. there is a low residual formate concentration in the reactor) (Fig. 2D). In this case, the electrochemical reaction (and therefore biomass productivity) is limited to ∼0.48 g L^-1^ hr^-1^ by the availability of CO_2_ at the electrode surface. Above ∼2.35 V, the steady-state concentration of CO_2_ at the cathode surface approaches zero (Fig. 2E), limiting the production rate of formate and therefore biomass productivity.

**Figure 2.**
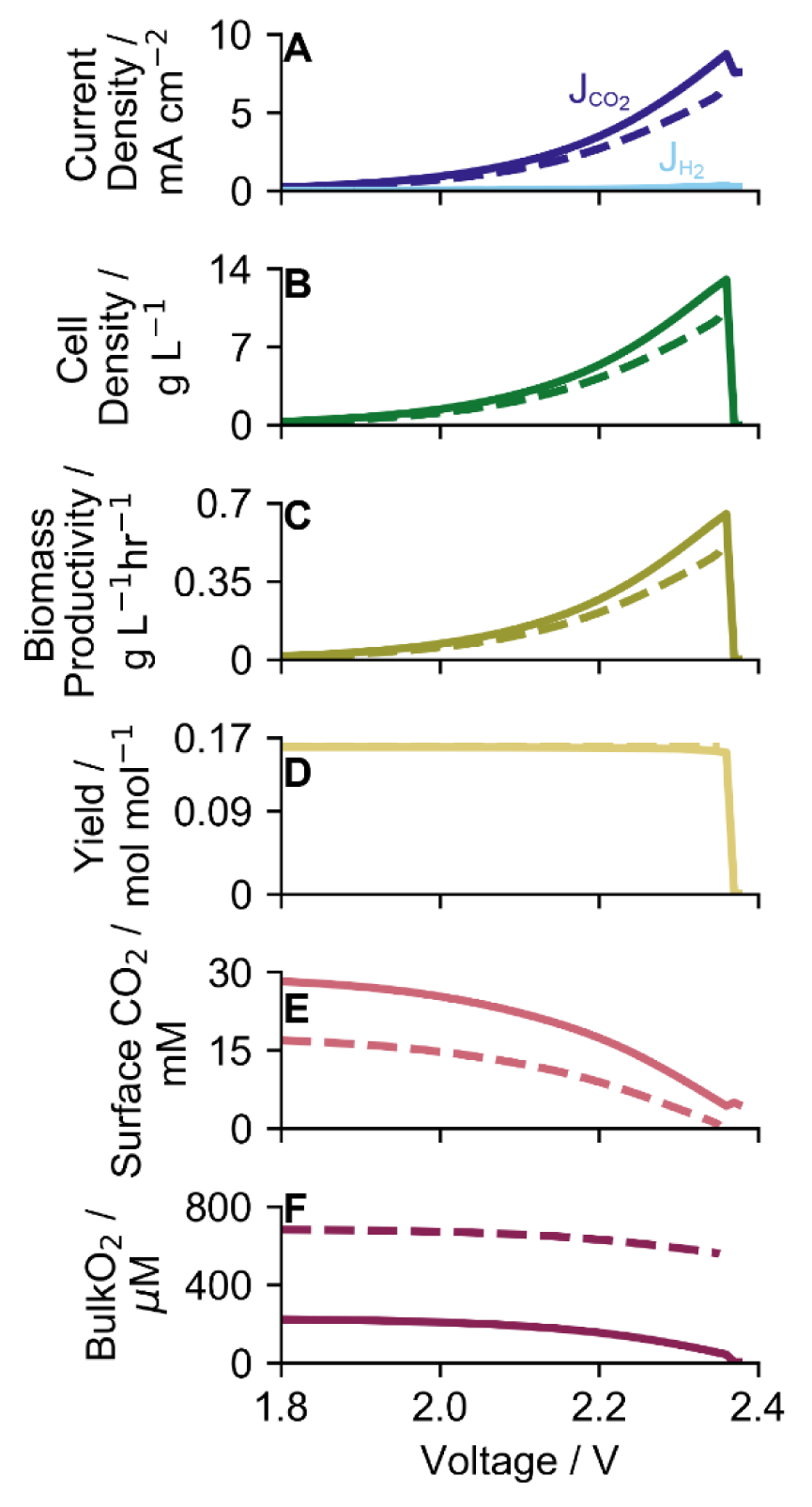
Current density and operating conditions as a function of applied voltage. Steady-state current densities towards CO_2_ (J_CO2_) and H_2_ (J_H2_), (**B**) cell density, (**C**) biomass productivity, (**D**) molar cell yield on formate, (**E**) CO_2_ liquid-phase concentration at the cathode surface, and (**F**) O_2_ liquid-phase concentration in the well-mixed bulk region as a function of applied voltage for 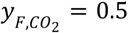 (dashed lines) and 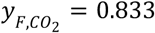 (solid lines) with S_A_ = 100 m^-1^ and D = 0.05 hr^-1^.

In contrast, for the case of 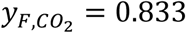, the steady-state concentration of CO_2_ at the cathode surface remains above 4 mM for the entire voltage range we consider (solid curve in Fig. 2E), but the cell density (Fig. 2B), productivity (Fig. 2C), and molar yield (Fig. 2D) all decrease rapidly to nearly zero above an applied voltage of ∼2.35 V. This behavior can be explained by microbial consumption of O_2_: above ∼2.35 V, the bulk O_2_ concentration drops below 5 μM (Fig. 2F), reducing the microbial growth rate (eqs. 8–9). When the growth rate is reduced, the residual formate concentration increases, reducing the yield (Fig. 2D). The combined effect of reduced growth rate and reduced cellular yield causes washout of the cells. Hence, the biomass productivity of the reactor in this case is limited to ∼0.65 g L^-1^ hr^-1^ by O_2_ transfer from the gas phase to the liquid phase. Interestingly, cell washout also reduces the electrode current density (Fig. 2A) because formate build-up reduces the pH in the reactor, increasing the Nernst potential drop at the anode surface and reducing OER kinetics because the OER exchange current density is pH-dependent (eq. 34, Table 1).

These results indicate that formate-mediated MES systems are limited either by CO_2_ or O_2_ transport and depend on the gas feed composition. When the O_2_ partial pressure is high, CO_2_ transport to the electrode surface limits formate production and therefore microbial growth; when the CO_2_ partial pressure is high, the O_2_ consumption rate driven by microbial respiration is limited by the gas/liquid mass transfer rate, slowing microbial growth and allowing build-up of HCOO^-^ to toxic concentrations. The trade-off between CO_2_ and O_2_ availability implies the existence of an optimal gas feed composition, which we explore next.

### 3.2 Reactor geometry determines optimal operating conditions and productivity

The optimal gas feed composition for biomass productivity is coupled to reactor design by the electrode surface area to reactor volume ratio (*S*_*A*_). This effect can be understood by considering the effective volumetric formate production rate (electrochemical CO_2_ consumption rate), which is proportional to the product of current density and *S*_*A*_ (*i*.*e*. 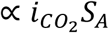). The current density 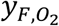 is limited by CO_2_ transport to the cathode surface. At equivalent current densities (*e*.*g*. the highest that can be supported by CO_2_ transport through the boundary layer), a larger *S*_*A*_ results in a larger volumetric formate production rate that must be matched by an increased microbial growth rate to maintain steady-state conditions. The increased microbial growth rate, then, results in a higher achievable cell density and volumetric productivity. However, microbial growth relies on O_2_ consumption (*via* respiration), so the O_2_ mass transfer rate from the gas phase to the liquid phase must also proportionally increase. If the k_L_a value is fixed, increasing the rate of O_2_ mass transfer can only be achieved by increasing the partial pressure of O_2_ in the gas feed, accomplished by increasing 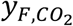 (equivalently, decreasing 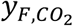).

To illustrate this effect, we calculated the maximum volumetric biomass productivities (g biomass per time per reactor volume) as a function of gas feed composition for different *S*_*A*_ values and as a function of *S*_*A*_ for different 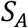 values (Fig. 3A, B). For an *S*_*A*_ of 100 m^-1^, the maximum volumetric productivity of ∼0.74 g L^-1^ hr^-1^ is reached when the gas feed is an 80/20 mixture of CO_2_/O_2_ (*i*.*e*. 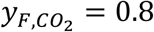) (Fig. 3A). CO_2_ transport to the cathode surface limits productivity when the CO_2_ gas fraction is <0.8, while CO_2_ gas fractions >0.8 result in O_2_ gas/liquid mass transfer-limited operation. Increasing *S*_*A*_ results in higher achievable volumetric productivities, reaching ∼1.76 g L^-1^ hr^-1^ with an *S*_*A*_ of 333 m^-1^ at 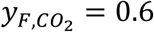. For a fixed gas composition, increasing *S*_*A*_ increases the volumetric productivity to a plateau value (∼0.74 g L^-1^ hr^-1^ for 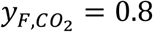), after which further increasing the *SA* has no effect (Fig. 3B). These plateau values correspond to the maximum microbial consumption rate that can be supported by O_2_ transfer from the gas phase to the liquid phase.

**Figure 3.**
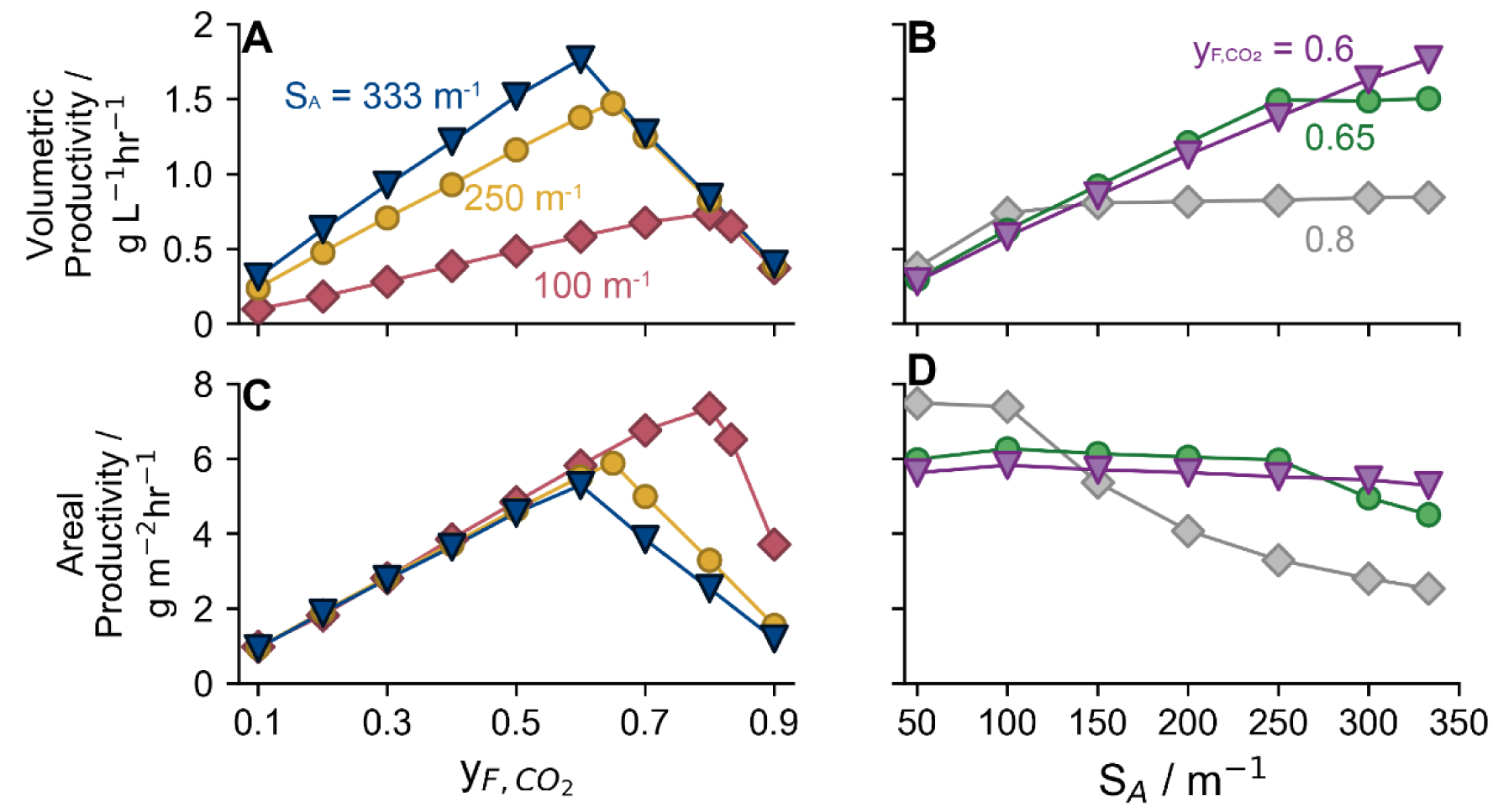
Coupled effects of gas feed composition and electrode surface area to volume ratio. Maximum volumetric productivity (**A, B**) and areal productivity (**C, D**), as a function of feed gas CO_2_ fraction 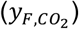 (**A, C**) and electrode surface area to volume ratio (*S*_*A*_) (**B, D**). Feed gas compositions plotted in (**B, D**) correspond to the optimal compositions for the three *S*_*A*_ values plotted in (**A, C**). All points use D = 0.05 hr^-1^.

Interestingly, higher areal biomass productivities (g biomass per time per cathode surface area) are achieved by decreasing *S*_*A*_ (Fig. 3C, D). This effect can also be understood by considering the volumetric production rate of formate. For a given gas composition, O_2_ gas/liquid mass transfer can support a maximal volumetric productivity (Fig. 3B), which corresponds to a specific volumetric production rate of formate (that is 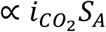). For a lower *S*_*A*_, a higher current density 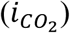 is necessary to achieve that rate. Because areal productivity is proportional to 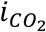, this results in an increased areal productivity as the *S*_*A*_ decreases (Fig. 3D). For 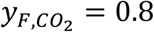, the areal biomass productivity plateaus at ∼7.4 g m^-2^ hr^-1^ for *S*_*A*_ <100 m^-1^, while 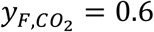 can only achieve ∼5.6 g m^-2^ hr^-1^ (a ∼25% decrease).

Results from this analysis have significant implications for mediated MES system engineering. First, the competing limitations of CO_2_ transport and O_2_ gas/liquid mass transfer place a fundamental upper bound on the productivity of mediated MES systems, and the optimal operating conditions (*i*.*e*. feed gas composition) are dependent on reactor design parameters (*S*_*A*_, *k*_*L*_*a*). The trade-off can be avoided by separating electrochemical formate production and biomass growth, as we discuss later, or by using gas diffusion electrodes (GDEs) to minimize the transport distance for CO_2_.^[54,59]^ Second, volumetric and areal productivities cannot be simultaneously optimized. A higher *S*_*A*_ results in higher volumetric productivity and therefore a higher CO_2_-fixing rate per unit volume, but also has a lower areal productivity and therefore requires more electrode material and is more resource-intensive. Hence, the optimal reactor design will be process-specific and will depend in part on the cell density or titer desired for a given product. Additionally, life-cycle assessments can inform reactor design and operation schemes that minimize energy and resource use while maximizing CO_2_-fixation.^[60,61]^

### 3.3 Decreasing dilution rate increases productivity

For a standard CSTR bioreactor where the growth substrate is fed in the liquid phase, the dilution rate can be adjusted to maximize biomass productivity.^[35]^ However, for a mediated MES system, the growth substrate is generated electrochemically, so substrate availability is partially decoupled from the dilution rate. To evaluate this effect, we calculated the biomass productivity as a function of dilution rate (Fig. 4). At a dilution rate of 0.01 hr, the volumetric productivity is ∼1.9 g L^-1^ hr^-1^ for 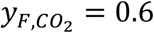 with an *S*_*A*_ of 333 m^-1^, but decreases monotonically as the dilution rate increases. For a simplified version of our model, where growth is dependent on a single, generic substrate and boundary layers are neglected, we show analytically that the productivity approaches a maximum as the dilution rate approaches 0, and that this result holds even if the yield decreases with increasing substrate concentration, as is the case for formate (Supplementary note 4).^[23]^ This result indicates that scaling-up the reactor volume offers an intrinsic benefit to mediated MES productivity. Increasing the reactor volume while maintaining a fixed volumetric flow rate will increase the overall productivity of the system and will result in a higher cell density (or product titer). However, a larger reactor also requires more electrode and other materials, increasing resource intensity, and more power for gas/liquid mass transfer (eq. 41). Life cycle analyses can indicate an optimal value for this trade-off and will be informed by the material and energy efficiency of the mediated MES system, which we consider in the next section.

**Figure 4.**
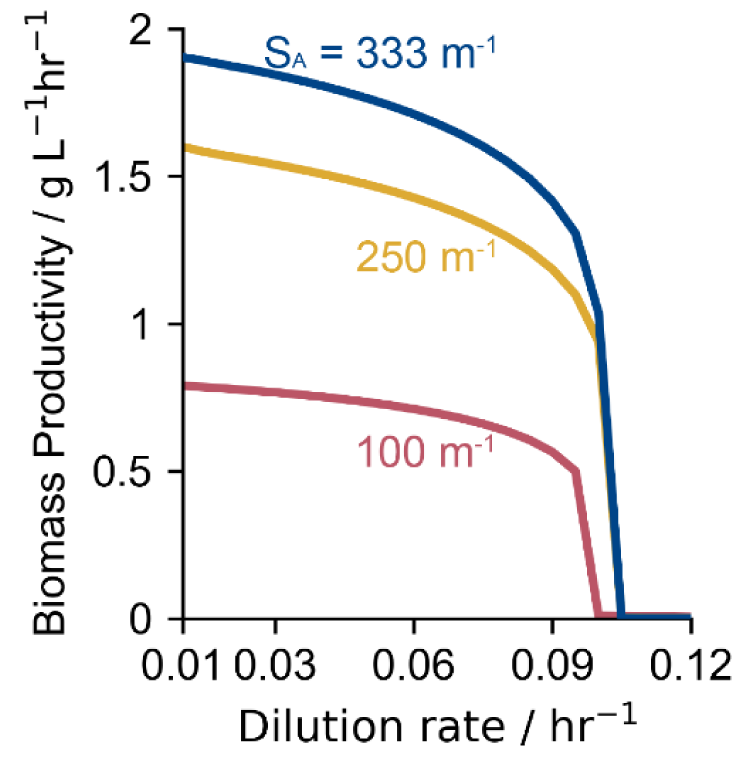
Effect of dilution rate. Steady-state volumetric biomass productivity as a function of the dilution rate for three different reactor operating configurations. Blue curve: S = 333 m^-1^, 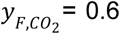. Yellow curve: S_A_ = 250 m^-1^, 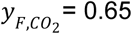. Red curve: S_A_ = 100 m^-1^, 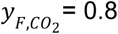.

### 3.4 Achievable carbon, hydrogen, and energy efficiency

Material utilization and energy efficiencies need to be defined and quantified as a function of reactor design and operating conditions because they will have a significant impact on the overall efficiency and practicality of MES systems. The carbon utilization efficiency can be written as the fraction of carbon exiting the reactor in biomass (since the total carbon fed to and exiting from the reactor must be equal):

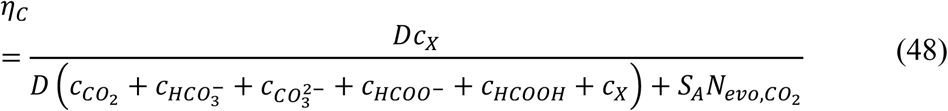

and the distribution of carbon exiting the reactor can be quantified simply by adjusting the numerator in this equation. For the formate-mediated MES system, <10% of the carbon fed to the reactor is diverted to biomass regardless of the applied potential (Fig. 5A). Notably, this is not due to low utilization of formate, which is nearly completely consumed prior to exiting the reactor. Instead, evolution of CO_2_ at the anode surface (eq. 42) comprises >80% of the carbon exiting the reactor. Gas recycle will therefore be necessary to achieve high overall carbon utilization efficiency for scaled MES systems.

**Figure 5.**
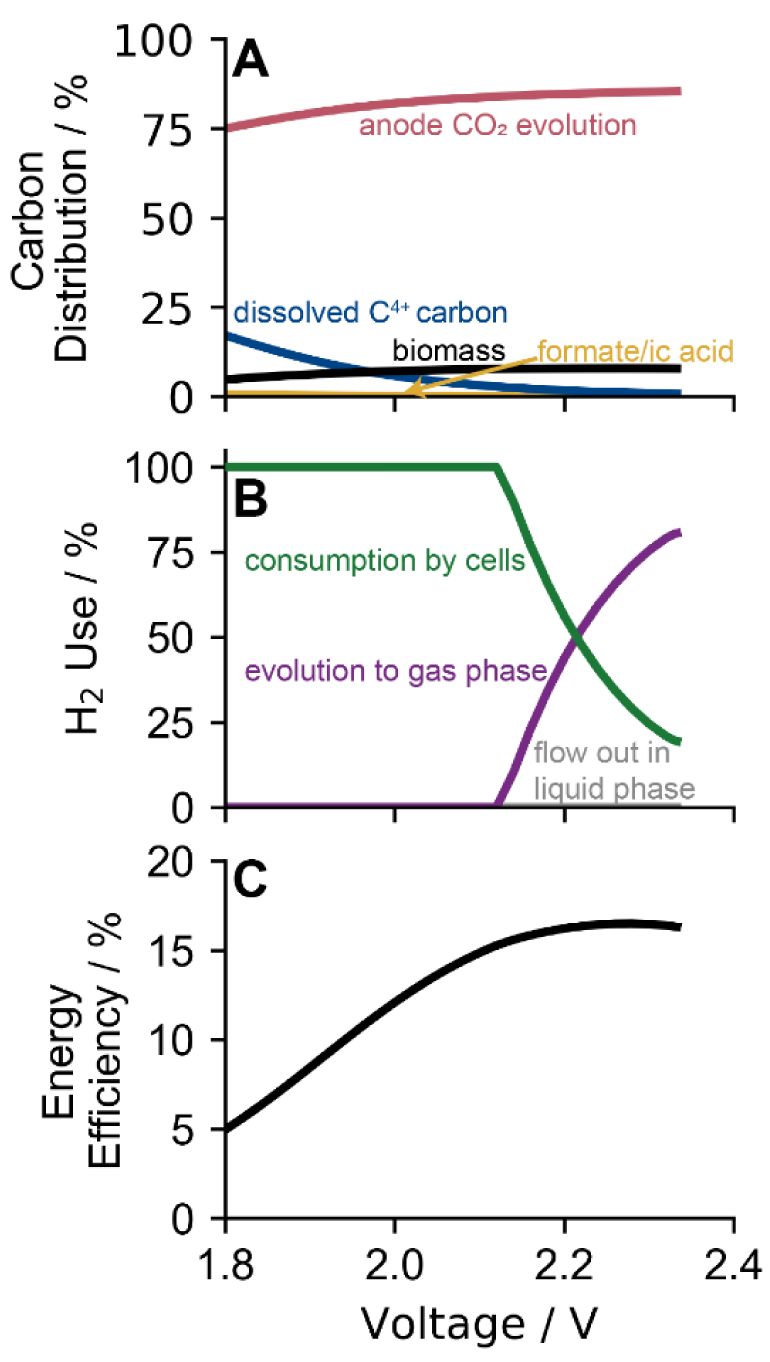
Carbon distribution, H_2_ utilization, and energy efficiency. (**A**) Distribution of carbon exiting the reactor; (**B**) final destination of H_2_ produced by the cathode; (**C**) energy efficiency towards cellular biomass as a function of applied voltage. All curves use D = 0.05 hr^-1^, S_A_ = 333 m^-1^, and 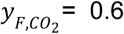. Dissolved C^4+^ carbon corresponds to solubilized CO_2_, HCO_3_ ^-^, and CO_3_^2-^.

H_2_ utilization efficiency can be calculated in a similar fashion, this time using the rate of H_2_ consumption by cells:

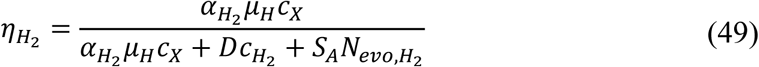

In our system, nearly all the H_2_ is consumed by cells at low applied voltages (Fig. 5B). However, as the voltage (and therefore, the current density) increases, evolution at the cathode surface begins to dominate because H_2_ is generated more rapidly than it can be solubilized by the liquid medium. This effect strongly limits the achievable biomass productivity for H_2_-mediated MES systems.

To calculate the energy efficiency, we used the estimate that autotrophic biomass production requires ∼479 kJ mol^-1^,^[6,62]^ and considered the power input from gas/liquid mass transfer and the applied current density:

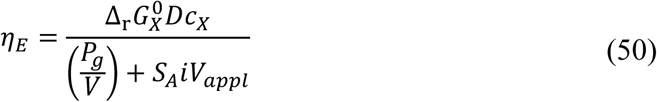

where 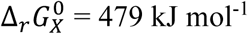 kJ mol^-1^ is the Gibbs’ free energy change for the biomass production reaction, *i* is the total current density and *V*_*appl*_ is the applied voltage. We note that this calculation should represent an upper bound on *η*_*E*_ since we do not consider the power demand for pumping liquid media and for heating or cooling the reactor to maintain optimal microbial growth temperatures. In our system, the energy efficiency increases from ∼5% at 1.8 V to a maximum of ∼16.5% at ∼2.3 V (Fig. 5C). Upon coupling to commercially available solar cells that have an energy efficiency of ∼20–25%, the system can therefore achieve an overall solar-to-chemical (STC) efficiency (*η*_*STC*_ = *η*_*solar*_ × *η*_*E*_) of only ∼3–4%.

Multiple strategies can overcome the low carbon utilization and energy efficiencies achievable for mediated MES systems. In addition to gas recycle, microbial engineering to improve growth yield can divert a higher fraction of carbon into biomass or products, reducing the futile CO_2_ cycle (CO_2_ reduced to formate electrochemically, then formate oxidized back to CO_2_ for metabolic energy by microbes), as we evaluate in the next section. Improved growth yield can also enhance overall energy efficiency, but energy efficiency may be better improved by separating electrochemical and microbial reactions into two reactors, allowing for individual, rather than coupled, optimization.

### 3.5 Engineered growth strategies can significantly improve productivity and efficiency

Although formate oxidation coupled to CO_2_ assimilation enables formatotrophic growth *via* the Calvin cycle, higher biomass yields are theoretically possible using the reductive glycine pathway (rGlyP; see Supplementary note 1 for additional details).^[24]^ We evaluated the potential for improved productivity, carbon utilization, and energy efficiency by formatotrophic growth using the rGlyP since this pathway has recently been engineered in *C. necator* and *E. coli*.^[21,22]^ The maximum biomass productivity for *E. coli* is ∼22% higher than for wild-type *C. necator* (*i*.*e. C. necator* using the Calvin cycle), reaching ∼2.15 g L^-1^ hr^-1^ at ∼2.3 V (Fig. 6A). Engineered *C. necator* slightly outperforms *E. coli*, reaching ∼2.23 g L^-1^ hr^-1^ at 2.33 V; the difference is attributable to the fact the *C. necator* is also able to use H_2_ as additional reducing power. Both the carbon utilization and energy efficiency are also significantly improved with the rGlyP, reaching ∼10.2% and ∼21.7% respectively for *E. coli*, representing ∼30% improvements over wild-type *C. necator* (Fig. 6B, C). Interestingly, the difference in operating temperature (30 °C for *C. necator* and 37 °C for *E. coli*) has almost no impact on productivity or efficiency limits despite impacting gas solubility (reduced with increased temperature), diffusivity of species (increased with increased temperature), acid/base equilibria (multiple effects), electrochemical potential (reduced with increased temperature), and electrochemical reaction kinetics (multiple effects), indicating that the competing impacts effectively cancel out. These results indicate the promise of microbial engineering to improve MES systems and can be combined with process engineering strategies to further increase productivity and efficiency.

**Figure 6.**
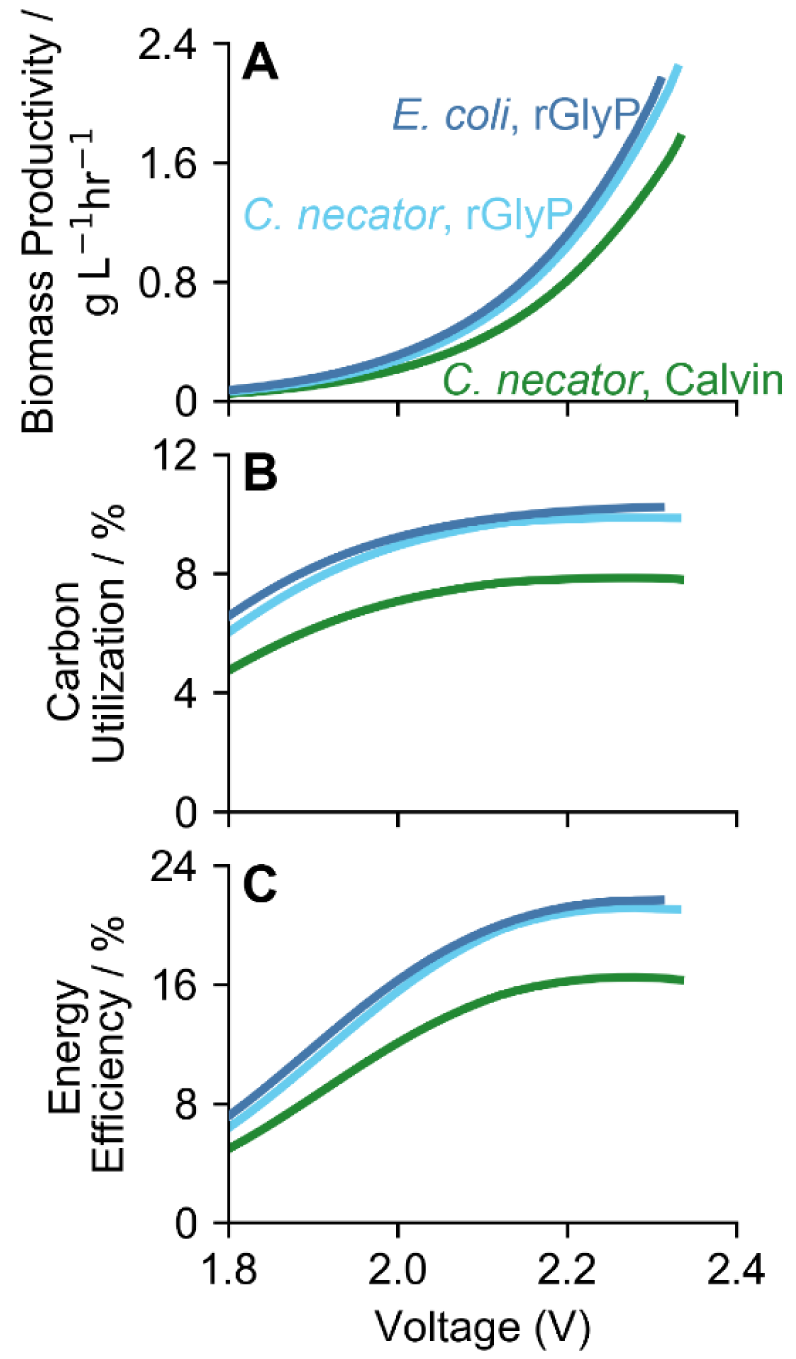
Effects of different formatotrophic growth strategies. (**A**) Productivity, (**B**) percent of fed carbon converted to biomass, and (**C**) energy efficiency towards cellular biomass for *C. necator* using the Calvin cycle (green curves) or the reductive glycine pathway (light blue curves) and *E. coli* using the reductive glycine pathway (dark blue curves) as a function of applied voltage. *C. necator* is modeled at 30°C; *E. coli* is modeled at 37 °C. All curves use S_A_ = 333 m^-1^ and 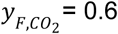.

### 3.6 Decoupled systems are likely to outcompete integrated reactors

Integrated systems for mediated MES minimize gas and fluid pumping power demand by integrating multiple processes into a single reactor. However, the competing requirements of CO_2_ and O_2_ mass transport create a fundamental limit on achievable productivity (Fig. 3). Decoupled systems, where an electrochemical reactor produces formate/ic acid at high rates that is then fed to a bioreactor (Fig. 7A), can break this limit by enabling individual optimization of the two processes. To evaluate the productivity of a decoupled system, we adapted our model by eliminating the electrochemical reactions, adjusting the gas feed composition to an 80/20 O_2_/CO_2_ ratio (*i*.*e*. 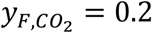), and altering the media feed to include sodium formate (HCOONa) and formic acid (HCOOH) in equilibrium at a pH of 2 to maintain reasonable Na^+^ concentrations of <3.5 g L^-1^ (note that the operating pH in the reactor remained at a pH of ∼7 due to microbial consumption of protons). We then calculated biomass productivity as a function of dilution rate and total inlet formate concentration, comprised of both HCOO^-^ and HCOOH (Fig. 7B). At lower inlet formate concentrations (<4.5 M), biomass productivity follows the standard trend of first increasing with increasing dilution rate up to a maximum value, then rapidly decreasing once microbial growth cannot match the dilution rate, causing washout. For this formate concentration range (<4.5 M), the maximum dilution rate that can be supported by microbes, and the achievable biomass productivity, also increase with inlet formate concentration following standard trends. However, above 4.5 M, microbial growth becomes O_2_-limited at higher dilution rates, causing a decline in the maximum dilution rate microbes can support. Despite the O_2_-limitation at high inlet formate concentrations, biomass productivities in excess of 2.4 g L^-1^ hr^-1^ are readily achieved with wild-type *C. necator*, outperforming the integrated MES system by >35% (Fig. 7B).

**Figure 7.**
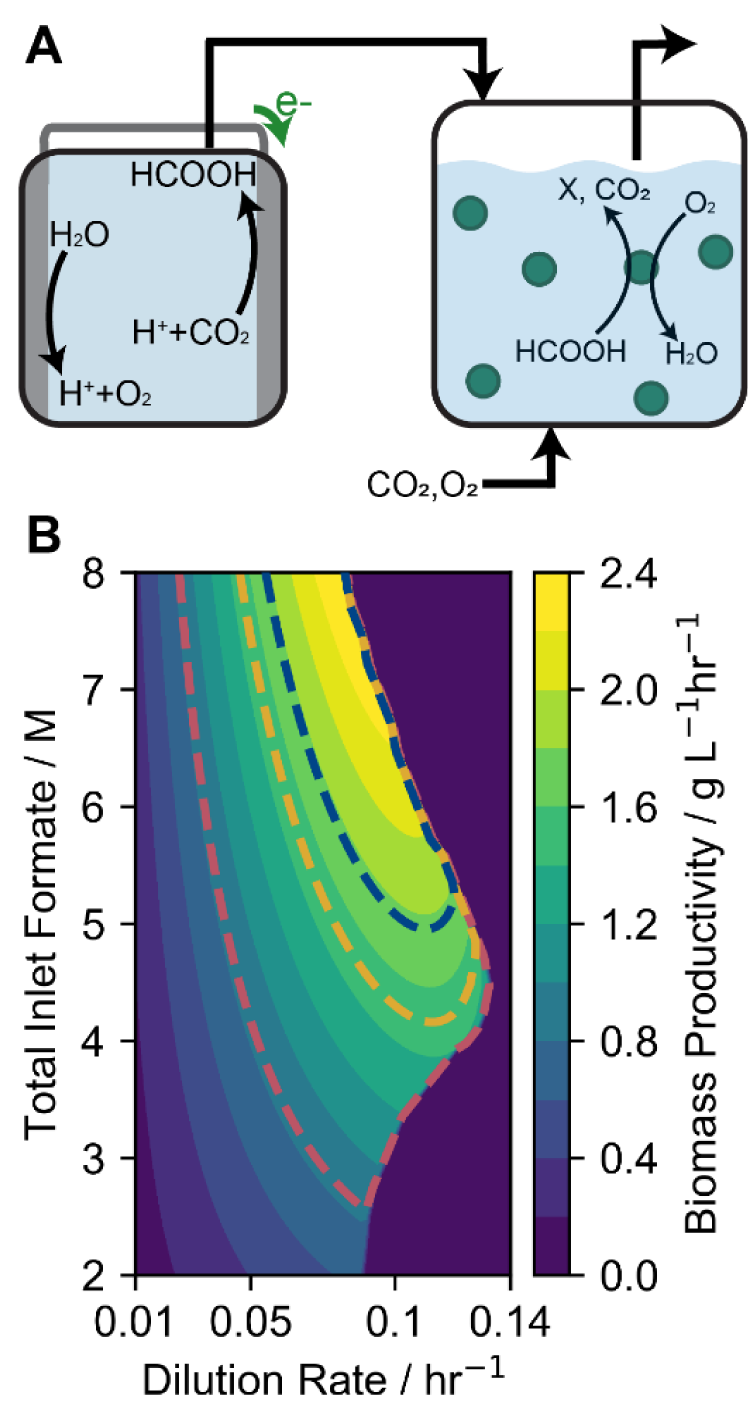
Higher productivity with decoupled systems. (**A**) Decoupled reactor scheme. Electrochemically-produced formate/ic acid is fed to a bioreactor along with CO_2_ and O_2_, which cells consume for growth. (**B**) Volumetric productivity as a function of dilution rate and total inlet formate concentration (comprised of sodium formate and formic acid) for a formate-fed bioreactor as described in the text with 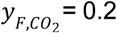. Overlaid dashed curves correspond to the maximum volumetric productivities of the integrated system reported in Fig. 3A (red: *S*_*A*_ = 100 m^-1^, yellow: 250 m^-1^, blue: 333 m^-1^).

A decoupled reactor system offers several benefits beyond higher biomass productivities. Higher applied current densities are achievable using GDEs or membrane-electrode assembly systems that can be optimized independently of the requirements for microbial growth (*i*.*e*. higher salt and buffer concentrations, higher pHs that enable higher selectivity towards carbon products).^[18,54,59]^ Issues associated with short-lived but toxic byproducts are lessened,^[12,28]^ but simple strategies to directly integrate electrochemical and microbial media with minimal intermediate processing are still necessary. The two-reactor system does have some challenges, however, including increased gas and liquid pumping requirements and the increased number of failure or contamination points. Life-cycle analyses comparing these reactor systems on the basis of material utilization, energy efficiency, and productivity should clarify these trade-offs and are enabled by the quantitative evaluation of mediated MES systems provided in this paper.

## 4. Conclusion

Formate-mediated MES represents a promising avenue for the production of multi-carbon molecules from CO_2_. In this study, we developed a comprehensive modeling framework for mediated MES systems that captures species transport, electrochemical, acid/base, and microbial reaction thermodynamics and kinetics, temperature effects, and gas/liquid mass transfer. We show that formate-mediated MES reactors are fundamentally limited by the trade-off between O_2_ gas/liquid mass transfer and CO_2_ transport to the cathode surface, and that decoupling electrochemical and microbial processes into separate reactors overcomes this limitation. We additionally evaluated the promise of synthetic formatotrophic growth via the reductive glycine pathway and showed that this strategy can significantly enhance carbon utilization and energy efficiency once the theoretical growth yields are realized. However, single-pass carbon utilization efficiency remains at ∼10% in the best case, indicating that gas recycle will be necessary for high overall CO_2_ utilization in scaled-up systems.

Future modeling efforts built on the framework we developed should include the effects of ionic strength on growth rate (especially for microbes such as *C. necator* that are sensitive to high salinity) and explicitly consider microbial product synthesis and its effects on growth yields and rates. Life-cycle assessments of different MES schemes (*i*.*e*. integrated vs. decoupled) should also be performed and can rely on the material and energy balances quantified here. The resulting optimal reactor design for mediated MES systems will enable a complete, rapid, and efficient process for the conversion of CO_2_ into a renewable chemical feedstock.

## Acknowledgements

This work was supported by the Center for the Utilization of Biological Engineering in Space (CUBES, https://cubes.space/), a NASA Space Technology Research Institute (grant number NNX17AJ31G). A.J.A. is supported by an NSF Graduate Research Fellowship under grant number DGE 1752814. We thank Dr. Jacob Hilzinger (UC Berkeley), Dr. Kyle Sander (UC Berkeley), and Jeremy Adams (UC Berkeley) for helpful discussions on microbial growth and model development, Helen Bergstrom (UC Berkeley) for useful discussions on electrochemistry, and Dr. Paul Tol (Netherlands Institute for Space Research, SRON) for a helpful reference on accessible color schemes (https://personal.sron.nl/~pault/).

## Author contributions

A.J.A. conceived of the idea, developed the model, analyzed data, and wrote the manuscript. D.S.C. edited the manuscript and supervised the project.

## Entry for the Table of Contents

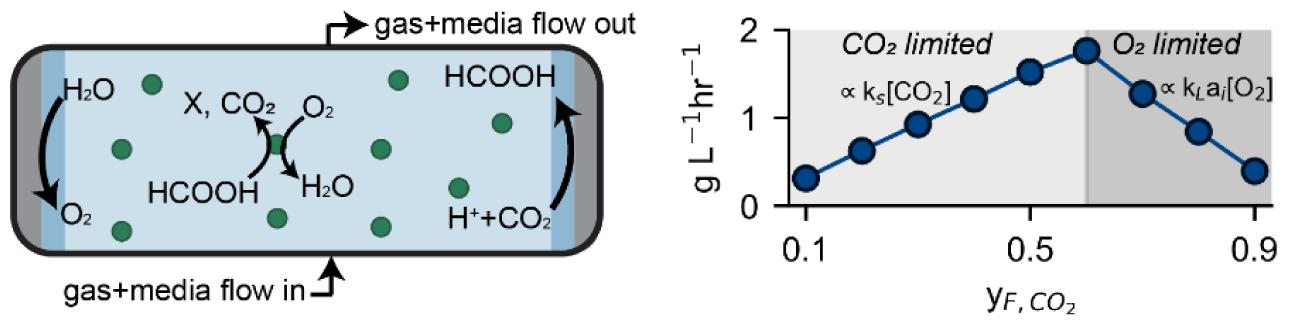

## Model-driven insight

Coupling electrochemical CO_2_ reduction to formate and microbial catalysis is a promising platform for sustainable chemical production. A comprehensive model capturing mass transport; electrochemical, acid/base, and microbial reaction kinetics and thermodynamics; temperature effects, and gas/liquid mass transfer was developed, revealing insights into performance limits for this system.

## References

[1] S. Perathoner, G. Centi, ChemSusChem. 2014, 7, 1274.

[2] N.J. Claassens, D.Z. Sousa, V.A.P.M. Dos Santos, W.M. De Vos, J. Van Der Oost, Nat. Rev. Microbiol. 2016, 14, 692.

[3] N.J. Claassens, I. Sánchez-Andrea, D.Z. Sousa, A. Bar-Even, Curr. Opin. Biotechnol. 2018, 50, 195.

[4] A. Prévoteau, J.M. Carvajal-Arroyo, R. Ganigué, K. Rabaey, Curr. Opin. Biotechnol. 2020, 62, 48.

[5] N.J. Claassens, C.A.R. Cotton, D. Kopljar, A. Bar-Even, Nat. Catal. 2019, 2, 437.

[6] D.G.N. Chong Liu, Brendan C. Colón Marika Ziesack, Pamela A. Silver, Science (80-.). 2016, 602.

[7] M. Soundararajan, R. Ledbetter, P. Kusuma, S. Zhen, P. Ludden, B. Bugbee, S.A. Ensign, L.C. Seefeldt, Front. Microbiol. 2019, 10,. https://doi.org/10.3389/fmicb.2019.01817.

[8] M. Wang, W. Zhong, S. Zhang, R. Liu, J. Xing, G. Zhang, J. Mater. Chem. A. 2018, 6, 9915.

[9] T. Krieg, A. Sydow, S. Faust, I. Huth, D. Holtmann, Angew. Chemie - Int. Ed. 2018, 57, 1879.

[10] J. Guan, S.A. Berlinger, X. Li, Z. Chao, V. Sousa e Silva, S. Banta, A.C. West, J. Biotechnol. 2017, 245, 21.

[11] W.O. Khunjar, A. Sahin, A.C. West, K. Chandran, S. Banta, PLoS One. 2012, 7, 44846.

[12] H. Li, P.H. Opgenorth, D.G. Wernick, S. Rogers, T. Wu, W. Higashide, P. Malati, Y. Huo, K.M. Cho, J.C. Liao, Science (80-.). 2012, 335, 1596.

[13] Y. Tashiro, S. Hirano, M.M. Matson, S. Atsumi, A. Kondo, Metab. Eng. 2018, 47, 211.

[14] M. Stockl, S. Harms, I. Dinges, S. Dimitrova, D. Holtmann, ChemSusChem. 2020,. https://doi.org/10.1002/cssc.202001235.

[15] R. Hegner, K. Neubert, C. Kroner, D. Holtmann, F. Harnisch, ChemSusChem. 2020, cssc. 202001272.

[16] T.D. Harrington, V.N. Tran, A. Mohamed, R. Renslow, S. Biria, L. Orfe, D.R. Call, H. Beyenal, Bioresour. Technol. 2015, 192, 689.

[17] T.D. Harrington, A. Mohamed, V.N. Tran, S. Biria, M. Gargouri, J.J. Park, D.R. Gang, H. Beyenal, Bioresour. Technol. 2015, 195, 57.

[18] Y. Chen, A. Vise, W.E. Klein, F.C. Cetinbas, D.J. Myers, W.A. Smith, T.G. Deutsch, K.C. Neyerlin, ACS Energy Lett. 2020, 5, 1825.

[19] R. Hegner, L.F.M. Rosa, F. Harnisch, Appl. Catal. B Environ. 2018, 238, 546.

[20] G. Wen, D.U. Lee, B. Ren, F.M. Hassan, G. Jiang, Z.P. Cano, J. Gostick, E. Croiset, Z. Bai, L. Yang, Z. Chen, Adv. Energy Mater. 2018, 8, 1.

[21] N.J. Claassens, G. Bordanaba-Florit, C.A.R. Cotton, A. De Maria, M. Finger-Bou, L. Friedeheim, N. Giner-Laguarda, M. Munar-Palmer, W. Newell, G. Scarinci, J. Verbunt, S.T. de Vries, S. Yilmaz, A. Bar-Even, BioRxiv. 2020, 2020.03.11.987487.

[22] S. Kim, S.N. Lindner, S. Aslan, O. Yishai, S. Wenk, K. Schann, A. Bar-Even, Nat. Chem. Biol. 2020, 16, 538.

[23] S. Grunwald, A. Mottet, E. Grousseau, J.K. Plassmeier, M.K. Popović, J.L. Uribelarrea, N. Gorret, S.E. Guillouet, A. Sinskey, Microb. Biotechnol. 2015, 8, 155.

[24] A. Bar-Even, E. Noor, A. Flamholz, R. Milo, Biochim. Biophys. Acta - Bioenerg. 2013, 1827, 1039.

[25] C.A. Cotton, N.J. Claassens, S. Benito-Vaquerizo, A. Bar-Even, Curr. Opin. Biotechnol. 2020, 62, 168.

[26] F. Enzmann, F. Mayer, M. Stöckl, K.M. Mangold, R. Hommel, D. Holtmann, Chem. Eng. Sci. 2019, 193, 133.

[27] L.F.M. Rosa, S. Hunger, T. Zschernitz, B. Strehlitz, F. Harnisch, Front. Energy Res. 2019, 7, 98.

[28] A. Sydow, T. Krieg, R. Ulber, D. Holtmann, Eng. Life Sci. 2017, 17, 781.

[29] S. Gadkari, M. Shemfe, J.A. Modestra, S.V. Mohan, J. Sadhukhan, Phys. Chem. Chem. Phys. 2019, 21, 10761.

[30] S. Gadkari, J.M. Fontmorin, E. Yu, J. Sadhukhan, Chem. Eng. J. 2020, 388, 124176.

[31] M. Kazemi, D. Biria, H. Rismani-Yazdi, Phys. Chem. Chem. Phys. 2015, 17, 12561.

[32] C. Picioreanu, K.P. Katuri, M.C.M. Van Loosdrecht, I.M. Head, K. Scott, J. Appl. Electrochem. 2010, 40, 151.

[33] C. Picioreanu, I.M. Head, K.P. Katuri, M.C.M. van Loosdrecht, K. Scott, Water Res. 2007, 41, 2921.

[34] C. Moß, N. Jarmatz, J. Heinze, S. Scholl, U. Schröder, ChemSusChem. 2020, cssc. 202001232.

[35] H.W. Blanch, D.S. Clark, Biochemical Engineering, 2nd ed., CRC Press, 1997.

[36] I.A. Figueroa, T.P. Barnum, P.Y. Somasekhar, C.I. Carlström, A.L. Engelbrektson, J.D. Coates, Proc. Natl. Acad. Sci. U. S. A. 2018, 115, E92.

[37] A. Ishizaki, K. Tanaka, J. Ferment. Bioeng. 1990, 69, 170.

[38] L. Rosso, J.R. Lobry, S. Bajard, J.P. Flandrois, Appl. Environ. Microbiol. 1995, 61, 610.

[39] J.M.T. Vasconcelos, S.C.P. Orvalho, A.M.A.F. Rodrigues, S.S. Alves, Ind. Eng. Chem. Res. 2000, 39, 203.

[40] L.A.H. Petersen, J. Villadsen, S.B. Jørgensen, K. V. Gernaey, Biotechnol. Bioeng. 2017, 114, 344.

[41] J.L. Meraz, K.L. Dubrawski, S.H. El Abbadi, K.H. Choo, C.S. Criddle, J. Environ. Eng. 2020, 146,. https://doi.org/10.1061/(ASCE)EE.1943-7870.0001703.

[42] P.M. Wilt, J. Colloid Interface Sci. 1986, 112, 530.

[43] M. Lin, L. Han, M.R. Singh, C. Xiang, ACS Appl. Energy Mater. 2019, 2, 5843.

[44] R.E. Zeebe, Geochim. Cosmochim. Acta. 2011, 75, 2483.

[45] D.E. Bidstrup, C.J. Geankoplis, J. Chem. Eng. Data. 1963, 8, 170.

[46] P. Han, D.M. Bartels, J. Phys. Chem. 1996, 100, 5597.

[47] D.R. Lide, ed., CRC Handbook of Chemistry and Physics, 84th edition, 84th ed., CRC Press, 2004. https://doi.org/10.1136/oem.53.7.504.

[48] K.G. Schulz, U. Riebesell, B. Rost, S. Thoms, R.E. Zeebe, Mar. Chem. 2006, 100, 53.

[49] P.W. Atkins, Physical Chemistry, 4th ed., Oxford University Press, 1990.

[50] Ulf Riebesell, Victoria J. Fabry, Lina Hansson, Jean-Pierre Gattuso, Guide to best practices for ocean acidification research and data reporting, 2011. https://doi.org/10.2777/66906.

[51] R.F. Weiss, Deep. Res. Oceanogr. Abstr. 1970, 17, 721.

[52] T.E. Crozier, S. Yamamoto1, Solubility of Hydrogen in Water, Seawater, and NaCI Solutions, 1974.

[53] H. Li, C. Oloman, J. Appl. Electrochem. 2007, 37, 1107.

[54] L.C. Weng, A.T. Bell, A.Z. Weber, Energy Environ. Sci. 2019, 12, 1950.

[55] H. Wang, D.Y.C. Leung, J. Xuan, Appl. Energy. 2013, 102, 1057.

[56] S. Haussener, C. Xiang, J.M. Spurgeon, S. Ardo, N.S. Lewis, A.Z. Weber, Energy Environ. Sci. 2012, 5, 9922.

[57] D.A. Stolpera, N.P. Revsbech, D.E. Canfield, Proc. Natl. Acad. Sci. U. S. A. 2010, 107, 18755.

[58] T. Takeshita, A. Ishizaki, J. Ferment. Bioeng. 1996, 81, 83.

[59] L.C. Weng, A.T. Bell, A.Z. Weber, Phys. Chem. Chem. Phys. 2018, 20, 16973.

[60] D. Helmdach, P. Yaseneva, P.K. Heer, A.M. Schweidtmann, A.A. Lapkin, ChemSusChem. 2017, 10, 3632.

[61] M. Shemfe, S. Gadkari, E. Yu, S. Rasul, K. Scott, I.M. Head, S. Gu, J. Sadhukhan, Bioresour. Technol. 2018, 255, 39.

[62] R.O.N. Grosz, G. Stephanopoulos, Biotechnol. Bioeng. 1983, 25, 2149.

