## Supplemental information for "Bioelectrochemical engineering analysis of formate-mediated microbial electrosynthesis"

### Supplementary information for Bioelectrochemical engineering analysis of formate-mediated microbial electrosynthesis

**Affiliations**
1. Department of Chemical and Biomolecular Engineering, University of California, Berkeley, CA 94720, USA
2. Molecular Biophysics and Integrated Bioimaging Division, Lawrence Berkeley National Laboratory, 1 Cyclotron Road, Berkeley, CA 94720, USA

**Supplementary note 1: Stoichiometric cell yield calculations using complete or partial formate oxidation or $H_2$ oxidation coupled to $CO_2$ and/or formate reduction**

*Formate oxidation coupled to $CO_2$ fixation*

Microbes support energy carrier (NAD(P)H and ATP) regeneration by using $NAD^+$-dependent formate dehydrogenases to catalyze the reaction

$$HCOO^- + NAD^+ \rightarrow CO_2 + NADH \qquad \text{(S1)}$$

NADH is then used to regenerate ATP following aerobic respiration (oxidative phosphorylation):

$$NADH + H^+ + \frac{1}{2}O_2 + (P/O)(ADP + P_i) \rightarrow NAD^+ + H_2O + (P/O)ATP \qquad \text{(S2)}$$

where $P/O$ is the oxidative phosphorylation ratio. We also assume NADH can be used to regenerate NAD(P)H according to:

$$1.33NADH + NADP^+ + 0.33H^+ \rightarrow 1.33NAD^+ + NADPH \qquad \text{(S3)}$$

If the Calvin cycle is used to fix $CO_2$, seven ATP and five NADH are consumed to fix three $CO_2$ molecules into one pyruvate molecule:

$$3CO_2 + 5NADH + 7ATP + 4H^+ \rightarrow C_3H_3O_3^- + 5NAD^+ + 7(ADP + P_i) + 3H_2O \qquad \text{(S4)}$$

The reductive glycine pathway makes more efficient use of energy carriers:

$$2HCOO^- + CO_2 + 2NADPH + 1NADH + 2ATP + 4H^+ \rightarrow C_3H_3O_3^- + 2NAD(P)^+ + 1NAD^+ + 2(ADP + P_i) + 3H_2O \qquad \text{(S5)}$$

in part because formate is a more energetic carbon source than $CO_2$. The resulting overall reaction for the production of pyruvate can then be written as

$$\left(5+\frac{7}{P/O}\right)HCOO^- + \left(4+\frac{7}{P/O}\right)H^+ + \frac{7}{2(P/O)}O_2 \rightarrow \left(2+\frac{7}{P/O}\right)CO_2 + \left(3+\frac{7}{P/O}\right)H_2O + C_3H_3O_3^- \quad \text{(S6)}$$

for the Calvin cycle, and as

$$\left(5.66+\frac{2}{P/O}\right)HCOO^- + \left(4.66+\frac{2}{P/O}\right)H^+ + \frac{1}{P/O}O_2 \rightarrow \left(2+\frac{2}{P/O}\right)CO_2 + \left(3.66+\frac{2}{P/O}\right)H_2O + C_3H_3O_3^- \quad \text{(S7)}$$

for the reductive glycine pathway.

From these equations, we calculate theoretical pyruvate yields on formate of 0.118 – 0.136 mol/mol using the Calvin cycle and 0.150 – 0.158 mol $mol^{-1}$ using the reductive glycine pathway for P/O ratios of 2 – 3. Therefore, to model the reductive glycine pathway, we increased the maximum experimental biomass yield on formate (0.169 mol $mol^{-1}$) by a factor of ~27% corresponding to the predicted increased pyruvate yield associated with a P/O ratio of 2.

*Hydrogen oxidation supporting $CO_2$ fixation*

NAD-reducing hydrogenases support energy carrier regeneration according to:

$$H_2 + NAD^+ \rightarrow H^+ + NADH \quad \text{(S8)}$$

Oxidative phosphorylation is used to regenerate ATP according to eq. (S2). Both the Calvin cycle and the reductive glycine pathway can be used to fix carbon. For the Calvin cycle, the overall equation for the production of pyruvate is given by

$$\left(5+\frac{7}{P/O}\right)H_2 + 3CO_2 + \frac{7}{2(P/O)}O_2 \rightarrow C_3H_3O_3^- + H^+ + \left(3+\frac{7}{P/O}\right)H_2O \quad \text{(S9)}$$

For the reductive glycine pathway, the overall equation is written as

$$\left(3.66+\frac{2}{P/O}\right)H_2 + CO_2 + 2HCOO^- + \frac{1.33}{P/O}O_2 + H^+ \rightarrow C_3H_3O_3^- + \left(3.66+\frac{2}{P/O}\right)H_2O \quad \text{(S10)}$$

From these equations, we calculate theoretical pyruvate yields on $H_2$ of 0.118 – 0.136 mol $mol^{-1}$ using the Calvin cycle and 0.215 – 0.231 mol $mol^{-1}$ using the reductive glycine pathway for P/O ratios of 2 – 3. To model the reductive glycine pathway in *C. necator*, we increased the experimental biomass yield on $H_2$ (~0.191 mol $mol^{-1}$) by a factor of ~82%, corresponding to the predicted increased pyruvate yield associated with a P/O ratio of 2. We also updated the biomassgenerating reaction to account for the additional consumption of formate and reduced consumption of $CO_2$.

**Supplementary note 2: boundary layer thickness**

The electrochemical reduction of $CO_2$ depends strongly on the concentration polarization that develops due to $CO_2$ consumption at the electrode surface.[1] Clark *et al.* measured the hydrodynamic boundary layer thickness in an aqueous cell as a function of the $CO_2$ gas flowrate (*i.e.* fluid mixing was generated *via* gas bubbling) and showed that increasing the flowrate decreased the boundary layer thickness but that the effect diminishes as the flowrate increased, approaching a minimum of ~40 µm at $CO_2$ flowrates >~20 sccm.[1] We assume a boundary layer of 100 µm (associated with a $CO_2$ flowrate of ~5 sccm in their system) in the main text and show how reduced boundary layer thicknesses impact biomass productivity in Fig. S1. The biomass productivity increases from ~1.5 g $L^{-1}$ $hr^{-1}$ to ~1.95 g $L^{-1}$ $hr^{-1}$ by decreasing the boundary layer thickness from 100 µm to 40 µm at an applied potential of 2.3 V (Fig. S1). Significant further enhancement is possible (to >2.5 g $L^{-1}$ $hr^{-1}$) by further reducing the boundary layer but this will likely require substantial agitation or other strategies to reduce the boundary layer thickness. Boundary layer thickness therefore plays a significant role in limiting the productivity of integrated MES systems and should be measured in experimental systems.

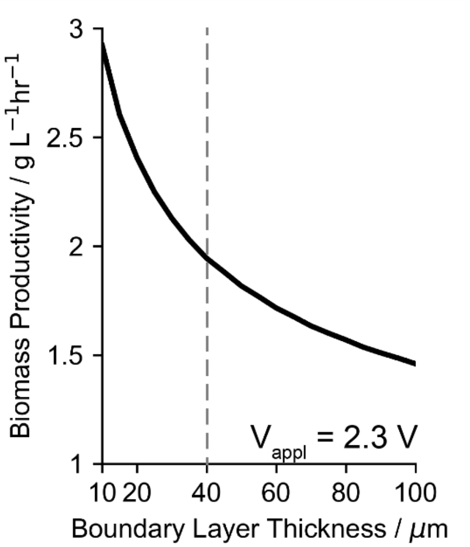


**Figure S1. Effect of boundary layer thickness on biomass productivity.** Biomass productivity at an applied voltage of 2.3 V as a function of the hydrodynamic boundary layer thicknesses for $y_{F,CO_2} = 0.6$ and an $S_A$ of 333 $m^{-1}$. Dashed line at 40 µm corresponds to the minimum boundary layer thickness Clark *et al.* (supplementary ref. 1) was able to achieve via fluid mixing due solely to gas bubbling.

**Supplementary note 3: diffusion coefficients of $Na^+$, $NO_3^-$, $HCOO^-$, $H^+$, $OH^-$, and $H_2$**

To estimate the temperature-dependence of the $Na^+$ diffusion coefficient, we used the well tabulated value of $1.334\times10^{-15}$ $m^2s^{-1}$ at 25 °C[2] and used this value to fit a Stokes radius ($R$) according to the Stokes-Einstein relationship:

$$R = \frac{k_B T}{6\pi D \mu} \quad \text{(S11)}$$

where $k_B$ is the Boltzmann constant and $\mu$ is the viscosity of water (in Pa·s). Once we solved for $R$, we used this value to determine the diffusion coefficient as a function of temperature by rearranging this same equation. We used the same procedure to determine the temperature-dependent diffusivity of $NO_3^-$ ions, this time using the value of $1.7\times10^{-9}$ $m^2s^{-1}$ at 25 °C.[3]

To describe the diffusivity of formate as a function of temperature, we used the Wilke-Chang correlation modified for carboxylic acids[4]:

$$D = \frac{6.6 \times 10^{-6} (XM)^{1/2} T}{\mu V^{0.6}} \quad \text{(S12)}$$

where $X = 2.6$ is the solvent constant for water, $M$ is the molecular weight of the solvent, and $V$ is the molar volume of the solute at normal boiling point.

To estimate the diffusion coefficients of $H^+$ and $OH^-$ in water, we used specific conductance data from Light *et al.*[5] and converted these values to diffusion coefficients following the Nernst-Planck equation, resulting in

$$D_i = \frac{RT\lambda_i}{z^2 F^2} \quad \text{(S13)}$$

where $\lambda_i$ is the specific conductance. We then fit these data to equations of the form:

$$D_i = Aexp\left[-B\left(\frac{1}{T} + \frac{1}{273.15}\right)\right] \quad \text{(S14)}$$

following Weng *et al*.[6] This equation fit $OH^-$ data well (Fig. S2A) but did not sufficiently capture $H^+$ data (Fig. S2B). For $H^+$, we attempted two additional fits, one following the functional form of $CO_2$, $HCO_3^-$, and $CO_3^{2-}$ diffusion coefficients (eq. S15) and a simple linear relationship (eq. S16):

$$D_i = A\left(\frac{T}{B} - 1\right)^C \quad \text{(S15)}$$

$$D_i = m(T - 273.15) + b \quad \text{(S16)}$$

The linear relationship fit best (Fig. S2B), so we used this relationship to describe the diffusivity of $H^+$ ions.

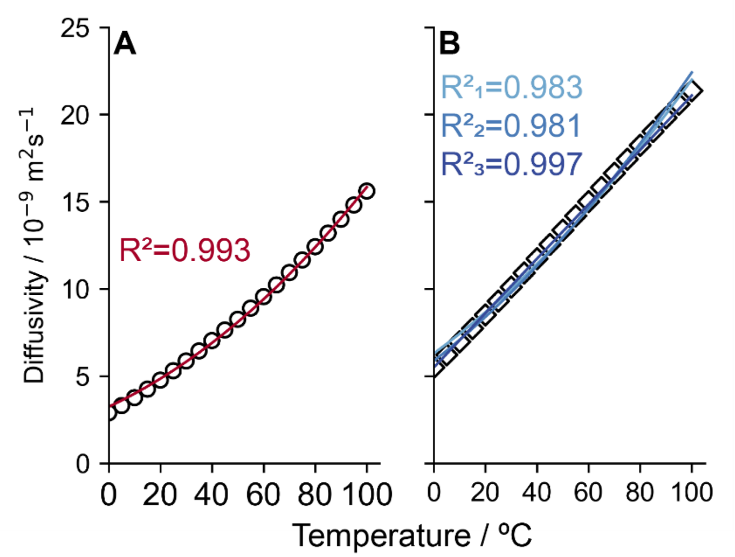


**Figure S2. Fitted diffusion coefficients of $OH^-$ and $H^+$ ions.** Experimental diffusion coefficients (black symbols) and fitted equations (colored, solid lines) for (**A**) $OH^-$ and (**B**) $H^+$ ions. Subscripts "1", "2", and "3" in (**B**) correspond to eqs. S14, S15, and S16 in the supplementary text.

For molecular hydrogen ($H_2$), the diffusivity was given as a function of temperature by Ferrell and Himmelblau[7] as:

$$D\ (cm^2 s^{-1}) = \frac{4.8 \times 10^{-7} T}{\mu^{\alpha}} \left(\frac{1 + \Lambda^{*2}}{V_m}\right)^{0.6} \quad \text{(S17)}$$

where $\mu$ is the viscosity of water (in centipoise), $V_m$ is the molar volume of $H_2$ at its normal boiling point, $\Lambda^*$ is the quantum parameter, and $\alpha$ is given by

$$\alpha = \frac{\sigma}{\left(\frac{V_m}{N}\right)^{1/3}} \quad \text{(S18)}$$

where $\sigma$ is the Lennard-Jones 6-12 potential. We used $V_m = 22.3897\ cm^3 mol^{-1}$ from Scott and Brickwedde[8], $\sigma = 2.96 \times 10^{-10}\ m$ and $\Lambda^* = 1.729$ from Nakanishi *et al.*,[9] and calculated $\mu$ (note that this equation returns $\mu$ in Pa·s, so the appropriate unit conversion to centipoise must be made) according to[10]

$$\log_{10} \mu = \frac{247.8}{T - 140} - 4.6173 \quad \text{(S19)}$$

Unfortunately, the resulting equation for $D$ did not sufficiently capture the data originally tabulated by Ferrell and Himmelblau, possibly because we used the viscosity of pure water. To account for this discrepancy, we adjusted the numerical fitting parameter, resulting in:

$$D = \frac{5.61 \times 10^{-7} T}{\mu^{\alpha}} \left( \frac{1 + \Lambda^{*2}}{V_m} \right)^{0.6} \tag{S20}$$

We simplified this equation to have only numerical parameters (except for the viscosity) in Table 1 in the main text. The original and improved fits are shown overlaid on the experimental data in Fig. S3.

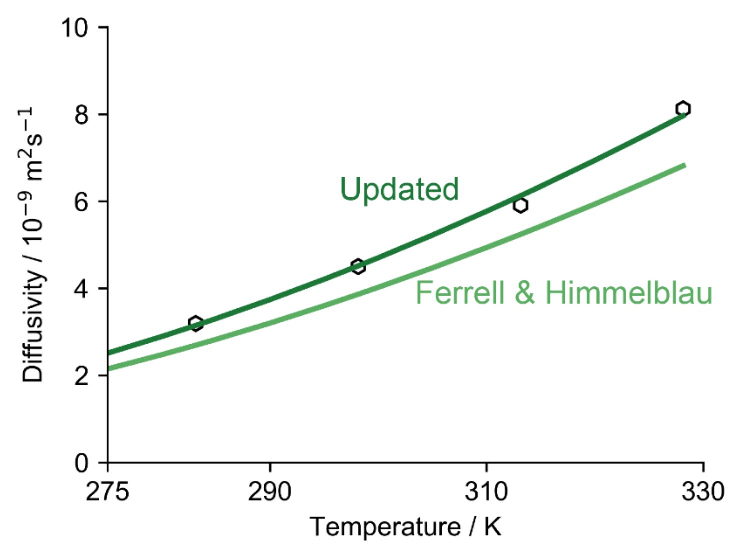


**Figure S3. Fitted diffusion coefficient of $H_2$.** Experimental diffusion coefficients (black symbols) and fitted equations (colored, solid lines) for $H_2$. The lighter solid line corresponds to the unmodified equation reported by Ferrell and Himmelblau (supplementary ref. 7); the darker solid line corresponds to the modification reported here.

**Supplementary note 4: derivation of the maximum productivity as a function of dilution rate for a simplified mediated MES reactor model**

For a simplified mediated MES reactor model, a generic substrate ($S$) is generated at an electrode surface and consumed by cells ($X$). The mass balances for substrate and cells are given as

$$\frac{dS}{dt} = -DS - \frac{1}{Y_{X/S}} \mu X + \frac{i S_A}{nF} \tag{S21}$$

and

$$\frac{dX}{dt} = \mu X - DX \tag{S22}$$

where $D$ is the dilution rate, $Y_{X/S}$ is the cell yield, $\mu$ is the specific growth rate, $i$ is the current density, $S_A$ is the electrode surface area to reactor volume ratio, $n$ is the number of electrons required to produce one $S$ molecule, and $F$ is Faraday's constant. Using standard Monod growth kinetics,

$$\mu = \mu_{max}\left(\frac{S}{K_S + S}\right) \tag{S23}$$

the steady-state substrate concentration and cell density are given by

$$S_{SS} = \frac{DK_S}{\mu_{max} - D} \tag{S24}$$

and

$$X_{SS} = -Y_{X/S}S_{SS} + \frac{Y_{X/S}iS_A}{DnF} \tag{S25}$$

The biomass productivity, $DX_{SS}$, is maximized with respect to the dilution rate by setting the derivative equal to 0:

$$\frac{d(DX_{SS})}{dD} = -Y_{X/S}S_{SS} = -Y_{X/S}\frac{DK_S}{\mu_{max} - D} = 0 \tag{S26}$$

which is satisfied only when $D = 0$. To confirm that this is a maximum, we confirm the second derivative is also <0:

$$\frac{d^2(DX_{SS})}{dD^2} = -Y_{X/S}\left[\frac{K_S\mu_{max}}{(\mu_{max} - D)^2}\right] < 0 \tag{S27}$$

The cell yield on formate is reduced as the formate concentration increases. For the generic substrate, we write this dependency as

$$Y_{X/S} = Y_{max}\left(1 - \frac{S_{SS}}{\theta_S}\right) \tag{S28}$$

and incorporate this into the derivative

$$\frac{d(DX_{SS})}{dD} = -Y_{X/S}S_{SS} = -Y_{max}S_{SS} + \frac{Y_{max}}{\theta_S}{S_{SS}}^2 = -Y_{max}S_{SS}\left[1 - \frac{S_{SS}}{\theta_S}\right] \tag{S29}$$

The maximum is then found by solving

$$-\frac{DK_S}{\mu_{max} - D}\left[1 - \frac{DK_S}{\theta_S(\mu_{max} - D)}\right] = 0 \tag{S30}$$

Again, $D = 0$ satisfies this equation, and the second derivative confirms that this corresponds to a maximum in the same way. We note that $D = \frac{\theta_S\mu_{max}}{K_S+\theta_S}$ also satisfies eq. (S30). However, this value corresponds to $S_{SS} = \theta_S$ and therefore $X_{SS} = 0$.

#### Supplementary references

[1] E.L. Clark, J. Resasco, A. Landers, J. Lin, L.T. Chung, A. Walton, C. Hahn, T.F. Jaramillo, A.T. Bell, *ACS Catal.* **2018**, *8*, 6560.

[2] D.R. Lide, ed., *CRC Handbook of Chemistry and Physics, 84th edition*, 84th ed., CRC Press, **2004**. https://doi.org/10.1136/oem.53.7.504.

[3] C. Picioreanu, M.C.M. Van Loosdrecht, J.J. Heijnen, *Water Sci. Technol.* **1997**, *36*, 147.

[4] D.E. Bidstrup, C.J. Geankoplis, *J. Chem. Eng. Data*. **1963**, *8*, 170.

[5] T.S. Light, S. Licht, A.C. Bevilacqua, K.R. Morash, *Electrochem. Solid-State Lett.* **2005**, *8*,. https://doi.org/10.1149/1.1836121.

[6] L.C. Weng, A.T. Bell, A.Z. Weber, *Phys. Chem. Chem. Phys.* **2018**, *20*, 16973.

[7] R.T. Ferrell, D.M. Himmelblau, *AIChE J.* **1967**, *13*, 702.

[8] R.B. Scott, F.G. Brickwedde, *J. Res. Natl. Bur. Stand. (1934).* **1937**, *19*, 237.

[9] K. Nakanishi, E.M. Voigt, J.H. Hildebrand, *J. Chem. Phys.* **1965**, *42*, 1860.

[10] J. Kestin, M. Sokolov, W.A. Wakeham, *J. Phys. Chem. Ref. Data*. **1978**, *7*, 101.
